# Single-cell profiling reveals stromal niche and α5β1 regulation of neutrophil phenotypes in ulcerative colitis

**DOI:** 10.64898/2026.09.21.753318

**Authors:** Yuxi Zhou, Jorge Canas, Jiayu Ye, Jacqueline N. Hoang, Sonia Ferkel-Soltau, Derek R. Holman, Raoul Sojwal, Touran Fardeen, Gustavo Vazques, Yuting Huang, Kathryn Peterson, Garry Nolan, Sidhartha Sinha, Stephan Rogalla, John Gubatan

**Author notes:** **Correspondence:** John Gubatan, MD. Division of Gastroenterology and Hepatology, Mayo Clinic Florida, 4500 San Pablo Road, Mangurian Building 5220, Jacksonville, FL 32224, USA. Yuxi Zhou and Jorge Canas share first authorship.

## Abstract

**Background and Aims:** Fibroblast–neutrophil interactions are implicated in ulcerative colitis (UC) treatment resistance, but whether fibroblasts modify neutrophil inflammatory activity remains unclear. We investigated responses to fibroblast exposure, stromal depletion, and α5β1-directed treatment.

**Methods:** We integrated human and mouse colonic single-cell RNA sequencing (scRNA-seq), spatial protein profiling (CODEX), fibroblast–neutrophil coculture, and dextran sulfate sodium (DSS) mouse colitis with fibroblast activation protein (FAP) ablation or α5β1-directed treatment. Readouts included cellular representation and protein fluorescence via flow cytometry, neutrophil extracellular trap (NET)-associated elastase activity, RNA programs and epithelial responses using scRNA-seq.

**Results:** Inflamed UC tissue contained higher fractions of inflammatory fibroblasts and oncostatin M (OSM)- and CXCR4-associated neutrophils. Inflammatory fibroblasts had higher FAP–α5β1 module scores, and proximity to FAP-high fibroblasts was associated with higher neutrophil OSM and CXCR4 fluorescence. UC fibroblasts increased neutrophil OSM, CXCR4 and myeloperoxidase (MPO) fluorescence and NET-associated elastase activity relative to control fibroblasts, while α5β1-directed treatment attenuated these responses. Both mouse FAP ablation and α5β1 blockade in DSS colitis reduced histologic inflammation and overall neutrophil frequency. Ablation broadly reduced recovered neutrophil representation, whereas blockade increased OSM-positive and PADI4-positive percentages within neutrophils despite lower MPO fluorescence within these subsets. Both interventions were associated with lower epithelial chemokine scores but distinct absorptive and mucus/secretory responses.

**Conclusions:** Fibroblast exposure modifies neutrophil inflammatory phenotype and effector-associated activity, identifying candidate therapeutic pathways in UC. Evaluating these responses alongside neutrophil representation and tissue inflammation could inform pharmacodynamic assessment of stromal-directed therapies.

**Graphical Abstract:** 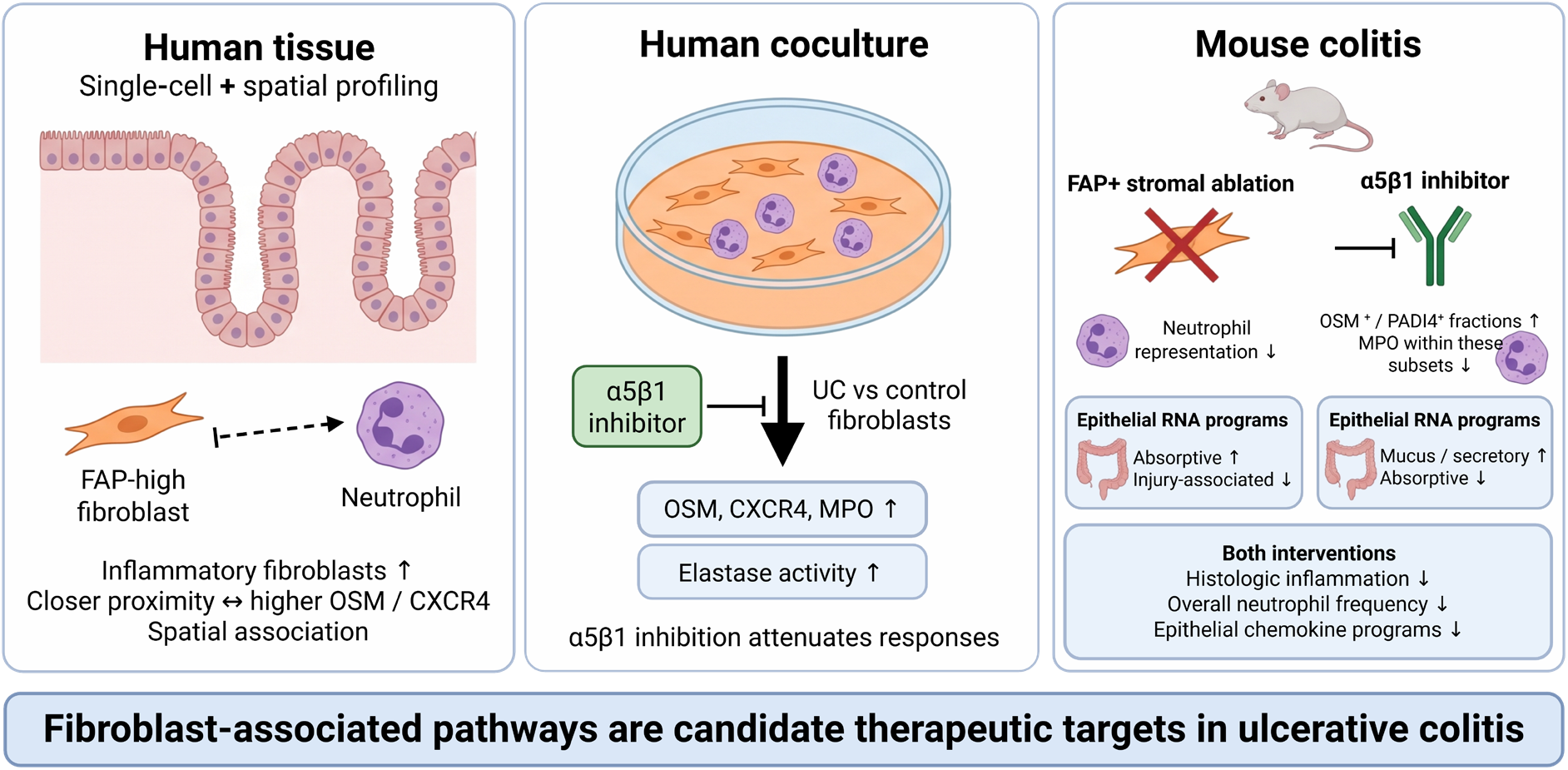

**What You Need to Know:** *Background and Context:* Fibroblast–neutrophil interactions are associated with treatment resistance in ulcerative colitis. Whether fibroblast exposure modifies neutrophil inflammatory phenotype and effector activity, beyond its established recruitment role, remains unclear.

*New Findings:* UC fibroblasts increased neutrophil inflammatory proteins and NET-associated elastase activity; α5β1-directed treatment attenuated these responses. FAP ablation and α5β1-directed treatment reduced colitis and overall neutrophil frequency. α5β1-directed treatment lowered MPO within OSM-positive and PADI4-positive neutrophils despite higher marker-positive percentages. Epithelial single-cell RNA sequencing revealed reduced chemokine programs with both interventions but distinct barrier-associated and regenerative expression patterns.

*Limitations:* Cellular drug targets remain unresolved. Spatial associations and elastase activity do not establish direct contact, NET formation, or clinical efficacy.

*Clinical Research Relevance:* Fibroblast-associated pathways offer candidate therapeutic opportunities to modify neutrophil inflammatory phenotype and effector activity in UC. Coculture responses were pharmacologically modifiable, and stromal interventions reduced experimental colitis. Measuring neutrophil frequency, subset proteins, and effector activity alongside tissue inflammation could guide pharmacodynamic assessment. Clinical efficacy and treatment-selection utility require validation.

*Basic Research Relevance:* Human coculture provides experimental evidence that fibroblast exposure modifies neutrophil inflammatory proteins and NET-associated elastase activity. FAP+ stromal ablation and α5β1-directed treatment produced distinct patterns of neutrophil representation and subset proteins in colitis. Epithelial single-cell RNA sequencing revealed reduced chemokine programs with both interventions but differing barrier-associated and regenerative expression patterns. Together, these findings motivate mechanistic studies of how stromal recruitment cues, soluble signals, and matrix-associated interactions regulate neutrophil responses, and whether these changes contribute to reduced epithelial inflammation and tissue repair.

*Lay Summary:* Fibroblasts, support cells in the intestine, increased inflammatory activity in immune cells called neutrophils. Blocking a pathway involved in cell attachment reduced these responses in cultures. Targeting fibroblast-associated pathways also reduced experimental colitis, with changes in intestinal lining gene activity related to inflammation, barrier function, and repair. These findings suggest potential approaches to treating ulcerative colitis.

## Introduction

Neutrophil-rich mucosal inflammation is a characteristic feature of active ulcerative colitis (UC).^1^ The consequences of this infiltration depend on the inflammatory properties of the cells present. Cytokines, chemotactic receptors, granule proteins, and metabolic programs capture different aspects of neutrophil biology, and single-cell studies have resolved maturation- and infection-associated diversity in mice.^2^ Neutrophil frequency alone therefore provides an incomplete description of mucosal inflammation. Defining the tissue signals that modify neutrophil phenotype and effector activity could identify therapeutic opportunities beyond reducing cell accumulation. Intestinal fibroblasts maintain extracellular matrix and epithelial niche signals^3^ and undergo substantial remodeling in inflammatory bowel disease (IBD). Inflammatory fibroblast populations and multicellular programs have been linked to treatment-resistant disease.^4,5,6^ Studies in stricturing Crohn’s disease further show how tissue profiling can nominate stromal adhesion molecules for functional investigation.^7^ These observations support examining fibroblasts as active regulators of the inflammatory environment.

Two established stromal–myeloid pathways inform this question. Oncostatin M (OSM) induces inflammatory and adhesion-related responses in intestinal stroma, and elevated mucosal OSM is associated with resistance to tumor necrosis factor (TNF) neutralization.^8^ Activated fibroblasts in intestinal ulcer beds also recruit neutrophils through interleukin-1 receptor–dependent programs associated with therapeutic nonresponse.^9^ These studies establish recruitment and cytokine responsiveness as important stromal functions, but leave open whether fibroblast exposure modifies the inflammatory properties of neutrophils already present.

Neutrophil heterogeneity makes this distinction clinically relevant. CD177-positive neutrophils in IBD exhibit enhanced bactericidal activity,^10^ whereas neutrophil extracellular traps (NETs) can sustain inflammatory signaling in UC.^11^ Our published UC atlas identified OSM-, CXCR4-, PADI4-, and MX1-associated neutrophil populations and linked state signatures and predicted stromal communication circuits to biologic therapy resistance.^12^ Those observations defined candidate cellular relationships, but marker-based annotation and communication inference did not establish how fibroblasts alter neutrophil protein phenotypes or effector activity.

We hypothesized that colonic fibroblast exposure modifies neutrophil inflammatory proteins and effector-associated activity, and that FAP depletion and α5β1-directed treatment produce different patterns of neutrophil representation and phenotype. We combined focused reanalysis of the published atlas with spatial protein profiling and scRNA-seq and flow cytometry of human fibroblast–neutrophil coculture and experimental mouse colitis. Tissue profiling established the disease context, coculture tested the influence of fibroblast exposure on neutrophil responses, and mouse interventions assessed accompanying changes in inflammation and epithelial programs. This design examines fibroblast regulation of neutrophil inflammatory properties alongside its established role in recruitment.

## Methods

### Study design and analytical populations

We reanalyzed our published human colonic single-cell RNA-sequencing atlas,^12^ available in the Broad Institute Single Cell Portal under accession SCP3755.^13^ The original cohort included 20 patients with moderate-to-severe UC, 18 with matched inflamed and noninflamed samples, and 408,930 cells including 47,516 neutrophils. Biopsies were processed within 2 hours; epithelial and lamina propria fractions were profiled separately using BD Rhapsody. Published quality control, Seurat processing, Harmony integration, and SCimilarity/manual annotation provided the source framework. Focused displays used 5,187 fibroblasts and 38,684 neutrophils. Eligibility yielded 13 fibroblast and 17 neutrophil biopsy pairs, 10 biopsies for receptor-expression comparisons, 14 paired patients for broader stromal programs, and 11 inflamed patients for matched neutrophil-state programs. CODEX included 24 distinct patients: 6 controls, 9 with noninflamed UC, and 9 with inflamed UC. Patients were the units for tissue-level comparisons; cells were not independent replicates. Supplementary Table 1 lists the CODEX antibodies; Supplementary Table 2 summarizes assay-specific populations and replication. The study was approved by Stanford University (IRB 28437, 60958, and 52317) for collection of blood and biopsies for co-culture experiments, scRNA-seq, and CODEX analyses.

### Discovery gene-program analyses

Fibroblast-associated programs were evaluated in the broader source stromal compartment from 14 paired UC patients. Neutrophil state programs used the same 11 inflamed UC patients with ≥20 cells in each of four annotated states. OSM, CXCR4, PADI4 and MX1 were excluded from module scoring wherever present; scores were averaged within patient-state and then equally across patients. Within-patient Friedman tests assessed state differences across nine programs. Supplementary Methods detail gene sets, scaling, annotation controls and cell-recovery sensitivity analyses.

### Multiplexed spatial protein profiling using CODEX

Co-detection by indexing (CODEX) used the PhenoCycler Fusion 2.0 system and established protocols.^14,15,16^ Oligonucleotide-conjugated antibody performance was assessed in tumor and tonsil controls and against the Human Protein Atlas.^17^ Following deparaffinization and pH-9 heat-induced epitope retrieval, sections were stained overnight at 4°C with 53 antibodies and DAPI (Supplementary Table 1), then fixed. The panel included FAP, α5β1, CD66b, CD15, CD16, CD11b, OSM, CXCR4, PADI4 and MX1. α5β1 was detected with clone M200 (Volociximab; Abcam, ab275977). Fluorescent reporters were imaged over successive cycles using PhenoImager Fusion software version 2.2. Registered, background-subtracted multichannel images were used for spatial analysis. Supplementary Methods provide staining, fixation, and imaging details.

Cell-level fluorescence and coordinates exported from QuPath^18^ were analyzed in Seurat 5.5.0^19^ using centered-log-ratio normalization, scaling, principal-component analysis and patient-level Harmony adjustment.^20^ Shared-nearest-neighbor Louvain clustering used Harmony dimensions 1–10 at resolution 1.0; UMAP used dimensions 1– 30. Cell labels combined lineage-marker profiles with selected markers and neutrophil-state definitions from our published UC atlas.^12^ CD66b, CD15, CD16 and CD11b supported neutrophil identity; OSM, CXCR4, PADI4 and MX1 guided phenotype annotation. Fibroblasts were assessed using CD140a, vimentin, podoplanin and FAP. Supplementary Methods detail the annotation workflow and retained analysis features.

### Spatial analysis

Six cellular neighborhoods summarized local cell-type composition within 50 μm, with equal patient weighting. Occupancy comparisons used exact patient-level permutation tests. Fibroblast proximity was evaluated against spatial nulls preserving local anatomy, density, and marker-category abundance. A separate joint model related neutrophil markers to continuous proximity to FAP-high, operationally α5β1-positive, epithelial, and endothelial reference populations. α5β1 positivity was defined by a posterior probability of at least 0.50 in the higher-signal component of a two-Gaussian mixture. Eight inflamed UC patients met model eligibility. This gate and the upper-quartile marker categories used for enrichment were analyzed separately.

### Coculture experiments and measurements

Human neutrophils were isolated from fresh whole blood obtained from non-IBD control participants using the MACSxpress® Whole Blood Neutrophil Isolation Kit, human (Miltenyi Biotec; cat. no. 130-104-434). This approach enriches unlabeled neutrophils through immunomagnetic depletion of other leukocytes and erythrocyte sedimentation. Fibroblasts were isolated separately from pooled control colonic biopsies and pooled inflamed UC colonic biopsies with a Mayo endoscopic subscore of 2, using Anti-Fibroblast MicroBeads, human (Miltenyi Biotec; cat. no. 130-050-601). Fibroblasts were seeded at 20,000 cells per well in six-well plates and expanded to approximately 75% confluence. Freshly isolated neutrophils were then added at 60,000 cells per well, and cocultures were maintained for 16 hours. Experimental conditions included neutrophils cultured alone, with control fibroblasts, or with UC fibroblasts. Treatment arms received the α5β1-directed antagonist ATN-161 at 20 µM (Tocris Bioscience; cat. no. 6058), the NAMPT inhibitor FK-866 at 10 µM (Tocris Bioscience; cat. no. 8072), or both agents at these concentrations. Inhibitor effects in neutrophils cultured without fibroblasts were evaluated separately from effects in coculture.

NET-associated neutrophil elastase activity was measured using the NETosis Assay Kit (Cayman Chemical; item no. 601010). Unbound elastase was removed by washing, the remaining extracellular material was digested with S7 nuclease, and released elastase activity was quantified with a chromogenic substrate at 405 nm. Figure 3D includes six independent experiments per group, measured on the same scale with the same normalization. This endpoint measures NET-associated elastase activity and does not independently establish NET formation.

Culture samples were analyzed by flow cytometry and BD Rhapsody single-cell RNA sequencing as described previously.^12^ Sample-median compensated neutrophil fluorescence was compared using exact unpaired permutation tests with Benjamini– Hochberg correction; elastase comparisons used exact unpaired permutation tests with Holm correction. RNA programs were summarized within recorded donor–condition samples containing at least 20 resolved neutrophils. Paired parametric tests and exact sign-flip sensitivity analyses were evaluated separately. Flow cytometry, elastase, and RNA assays were not assumed to use matched samples; independent elastase replication does not establish matching or independence for the other assays.

### Mouse colitis experiments and tissue analyses

FAP-TK mice (C.B6-Tg(Fap-TK)MRkl/J; The Jackson Laboratory, stock no. 035907) and wild-type mice received 2.5% dextran sulfate sodium (DSS) for five days and were euthanized on day 6. FAP-TK mice received ganciclovir intraperitoneally at 100 mg/kg per dose twice daily for five days or phosphate-buffered saline (PBS). Wild-type DSS mice received ATN-161 intraperitoneally at 100 mg/kg on days 1 and 3 or PBS. These were separate intervention experiments, not a randomized head-to-head comparison. Harvested colons underwent histologic scoring by a blinded pathologist using published criteria^21^, flow cytometry, and BD Rhapsody profiling. Flow cytometry distinguished marker-positive percentages within neutrophils from MPO fluorescence within OSM-, PADI4- and CXCR4-positive gates (Figure 5B; Supplementary Figure 10C–G). Histology and flow cytometry retain assay-specific statistical annotations detailed in Supplementary Methods. Milo summaries retain the original FDR calls and dominant state labels.^22^ Epithelial counts were aggregated by biological mouse, normalized using the trimmed mean of M-values,^23^ and expressed as log₂ counts per million for scoring. Treatment and acquisition batch were aligned in the mouse transcriptomic datasets; transcriptomic comparisons therefore describe intervention-associated patterns inseparable from potential batch effects. Animal procedures were approved by Stanford University (IACUC protocols 22000 and 27715) and Mayo Clinic (A00008640-26).

Epithelial programs were assessed across the whole compartment and within states meeting prespecified mouse and cell-count thresholds. Gene-level displays used stored mature-colonocyte log₂ CPM values, standardized across eligible mice for visualization; the complete 39-gene display accompanies the selected-gene panel. Individual-mouse plots combined existing chemokine, absorptive and mucus/secretory scores without new hypothesis tests. Supplementary Methods detail gene membership, eligibility and display transformations.

### Communication and external human analyses

NicheNet and MultiNicheNet nominated expression-based ligand–target relationships; priorities indicate regulatory potential rather than signaling flux.^24,25^ Independent human analyses used the TAURUS UC subset^26,27^ and paired spatial sections from one SCP3818 donor.^28^ TAURUS receptor analyses averaged activated-versus-other fibroblast differences within biopsies and then within 16 patients contributing 50 biopsies. TAURUS retained no neutrophils; its myeloid summary is a monocyte proxy. These datasets evaluate the stromal context and receptor-component expression, rather than replicate the complete experimental mechanism.

### Statistical analysis

Pairwise tests were two-sided, with endpoint-specific Benjamini–Hochberg (BH) correction unless Holm correction or assay-specific source annotations were specified. Discovery families comprised eight paired stromal tests, nine within-patient Friedman neutrophil-state tests, and five paired coexpression routes. Patients or biological mice were the analysis units where established; culture analyses used recorded sample or donor units. Neutrophil frequency denotes a proportion within the assay-specific parent population, marker-positive percentage denotes positivity within the neutrophil gate, and subset MPO denotes fluorescence within the specified marker-positive gate. RNA-state fractions use the stated recovered-cell compartment as denominator. These measurements do not establish absolute cell numbers. Intervals were pointwise; overlapping Milo neighborhoods were not independent state-level tests. Nonsignificance was not interpreted as equivalence. Supplementary Methods specify correction families, eligibility, and unresolved assay metadata.

## Results

### Inflamed UC tissue is enriched for inflammatory fibroblasts and selected neutrophil states

We first defined the human tissue context for fibroblast regulation of neutrophils. Five fibroblast states and OSM-, CXCR4-, MX1-, PADI4-, and low-RNA neutrophil populations were resolved in the discovery atlas (Figure 1A). In paired UC biopsies, inflammatory fibroblasts increased by a median 28.3 percentage points within the fibroblast compartment (13 pairs; q = .00715), whereas lamina propria and crypt-bottom fibroblast fractions decreased. Within the neutrophil compartment, OSM- and CXCR4- associated state fractions increased by 10.6 and 10.5 percentage points (17 pairs; q = .00585 and .00715). MX1- and PADI4-associated changes did not pass correction (Figure 1B).

**Figure 1.**
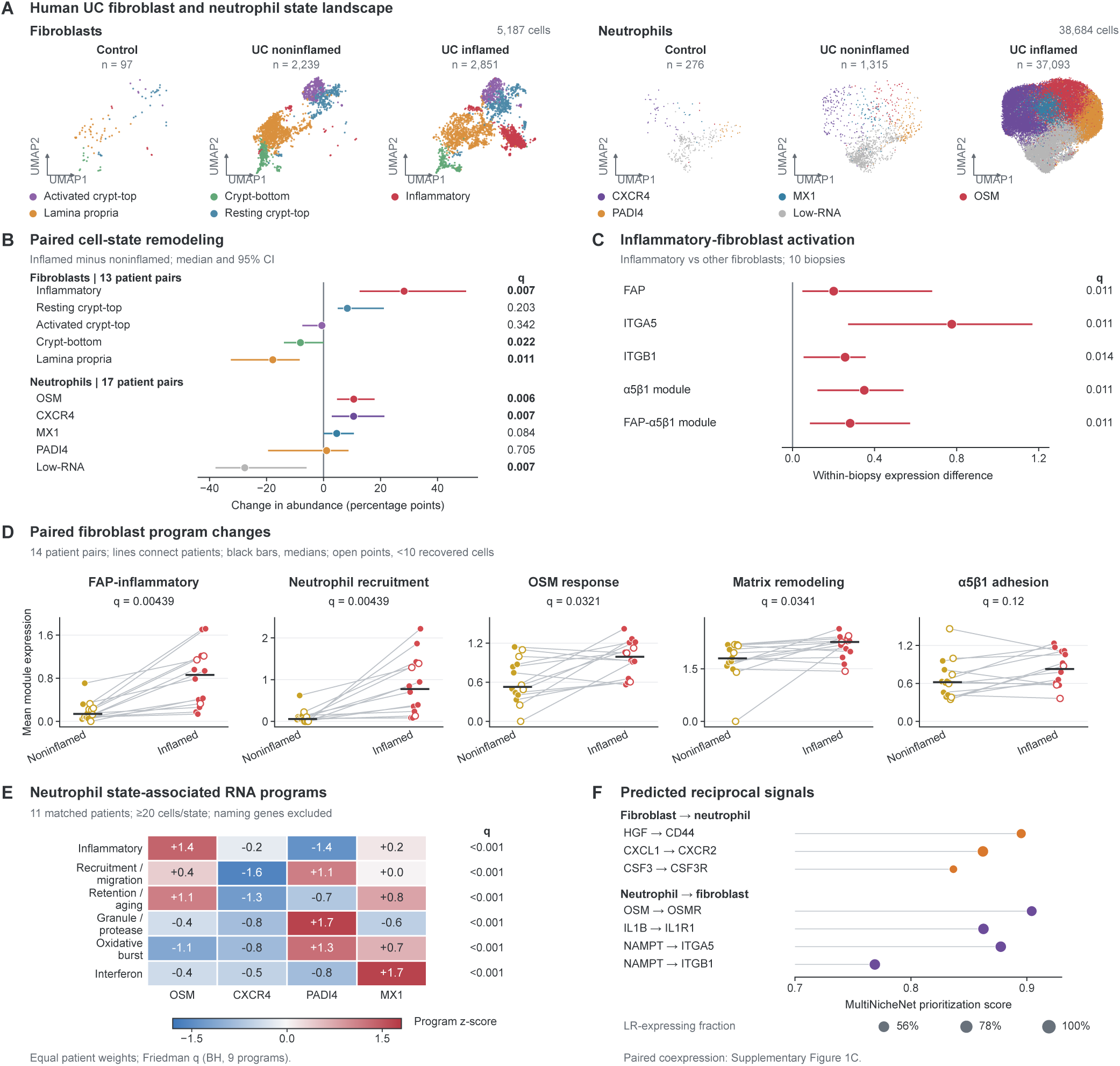
Inflammatory stromal programs and heterogeneous neutrophil states in human UC. (A) Fibroblast (5,187 cells) and neutrophil (38,684 cells) UMAPs. (B) Paired inflamed-minus-noninflamed state-fraction differences within each lineage; 13 fibroblast and 17 neutrophil patient pairs. (C) Inflammatory-minus-other fibroblast expression differences within 10 biopsies. B shows median paired differences; B,C show bootstrap 95% confidence intervals (CIs) and paired Wilcoxon tests with Benjamini–Hochberg (BH) correction; C covers five genes/programs. (D) Five fibroblast-associated programs in the broader stromal compartment, including smooth-muscle cells and pericytes; 14 patient pairs. Lines connect samples, bars indicate medians, and open points mark <10 cells. Wilcoxon tests are BH-adjusted across eight programs. (E) Marker-excluded neutrophil programs in 11 matched inflamed UC patients with ≥20 cells per state. Colors indicate z-scores across four equally patient-weighted state means. Friedman omnibus tests are BH-adjusted across nine programs; they do not establish pairwise differences. (F) Selected predicted communication priorities; dot size indicates expressing-cell fraction. NAMPT–ITGA5 and NAMPT–ITGB1 are separate component scores. Supplementary Figure 1 contains annotation controls and paired coexpression. Patients are replicates; composition, sparse recovery, and prediction limits are detailed in Supplementary Methods.

Inflammatory fibroblasts had higher FAP, ITGA5 and ITGB1 expression and α5β1-associated module scores than other fibroblasts within 10 biopsies (q = .0114–.0144; Figure 1C). Across the broader stromal compartment in 14 paired patients, FAP-inflammatory, neutrophil-recruitment, OSM-response and matrix-remodeling scores increased (q = .00439–.0341), whereas the α5β1-adhesion program did not pass correction (q = .120; Figure 1D; Supplementary Figure 1A). These compartment-level differences can reflect both cellular composition and expression. Requiring at least five recovered cells per condition retained nine pairs and significance for FAP-inflammatory and recruitment programs; at least 10 cells retained six pairs and no BH-significant program.

Neutrophil states also differed in their RNA profiles within the same 11 inflamed UC patients after excluding the four state-naming genes from module scoring. The OSM-associated state had the highest mean inflammatory score; the PADI4-associated state had the highest mean recruitment, granule/protease and oxidative-burst scores; and the MX1-associated state had the highest mean interferon score (Figure 1E). Each displayed program showed an omnibus state difference (all q < .001; BH correction across nine programs), without identifying specific significant pairwise contrasts. Original-score and cell-count sensitivity analyses retained these leading profiles, although retention/aging rankings depended on marker inclusion. The pooled annotation-control display uses a different denominator and retains the naming genes (Supplementary Figure 1B). These findings describe state-associated transcriptional heterogeneity rather than measured effector activity.

Communication inference nominated recruitment, matrix/adhesion, and reciprocal cytokine routes, including CXCL1–CXCR2, CSF3–CSF3R, OSM–OSMR, and IL1B–IL1R1 (Figure 1F). The three displayed paired coexpression comparisons did not pass correction (nine pairs; q = .055–.076; Supplementary Figure 1C). The coexistence of inflammatory stromal programs and heterogeneous neutrophil states prompted us to examine their spatial relationships in human tissue.

### FAP-high fibroblast proximity is associated with selected neutrophil proteins in UC

CODEX profiling identified 245,020 cells, including 14,906 neutrophils, across 24 independent patients (Figure 2A; Supplementary Figure 2). Six neighborhoods captured differences in local cellular composition (Figure 2B). CN3 tissue occupancy differed between inflamed UC and control (q = .0135) and between inflamed and noninflamed UC (q = .0347; Figure 2C), demonstrating patient-level remodeling of mucosal organization.

**Figure 2.**
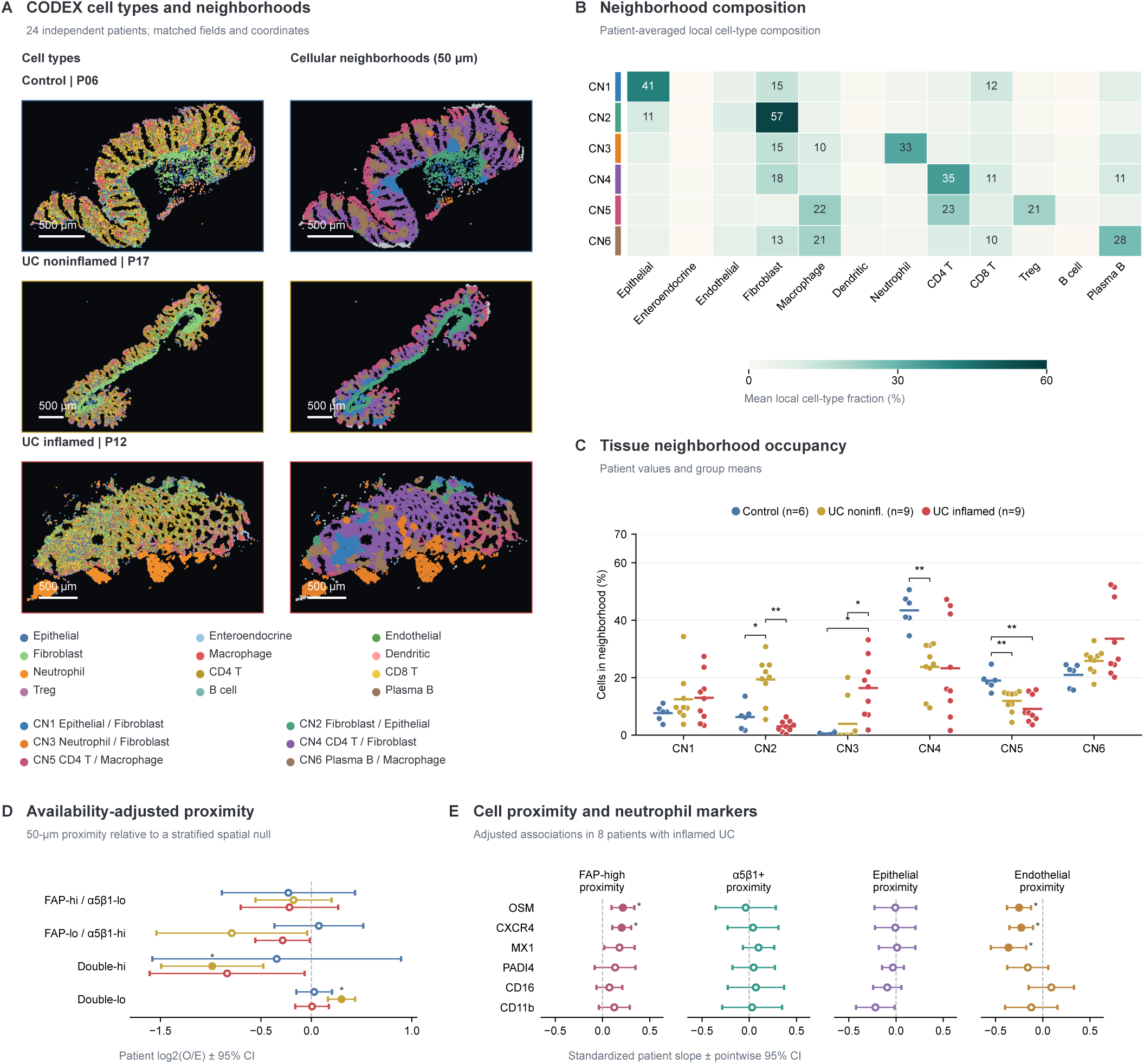
Cellular neighborhoods and fibroblast proximity in UC. (A) Representative cell-type and neighborhood maps; control P06, noninflamed UC P17, and inflamed UC P12. Gray indicates excluded assignments; source-coordinate scale bars, 500 μm. The cohort includes 6 control, 9 noninflamed UC, and 9 inflamed UC patients. (B) Patient-weighted cell composition in six 50-μm neighborhoods. (C) Tissue occupancy; lines indicate means. Exact patient-level permutation tests are BH-adjusted across 36 occupancy comparisons. (D) Availability-adjusted neutrophil– fibroblast proximity using upper-quartile FAP/α5β1 categories; mean patient log₂ observed/expected enrichment and 95% t CIs. Group sizes are 6/9/9, except double-high (5/9/7); BH correction covers 24 comparisons. (E) Adjusted marker–proximity coefficients in eight inflamed UC patients; means and 95% t CIs. α5β1 positivity uses mixture posterior ≥0.50; FAP-high uses the upper quartile. Models adjust jointly for reference-cell distances and local density; BH correction covers 24 tests. Filled symbols indicate q<0.05. *q<0.05; **q<0.01. Proximity does not establish direct signaling.

Availability-adjusted neutrophil proximity differed between inflamed and noninflamed UC for the double-low fibroblast category (q = .0240; Figure 2D). In eight eligible inflamed UC patients, proximity to FAP-high fibroblasts was positively associated with neutrophil OSM and CXCR4 fluorescence after joint adjustment for other reference-cell distances and local density (both q = .0191; Figure 2E). None of the six associations with proximity to operationally α5β1-positive fibroblasts passed correction; the mixture-based gate selected 806 of 44,516 fibroblasts (Supplementary Figure 3). The FAP-high associations linked stromal location to neutrophil protein phenotype, motivating experimental testing of fibroblast exposure in coculture.

### UC fibroblasts increase neutrophil inflammatory proteins and NET-associated elastase activity

To test whether fibroblast exposure could modify neutrophil responses outside inflamed tissue, we cocultured non-IBD control-blood neutrophils with control or UC fibroblasts (Figure 3A). UC fibroblasts increased sample-median neutrophil OSM, CXCR4, and MPO fluorescence relative to control fibroblasts (all q = .00928). ATN-161, an α5β1 inhibitor, within UC coculture reduced OSM and CXCR4 (both q = .00749) and MPO fluorescence (q = .0127; Figure 3B and C).

**Figure 3.**
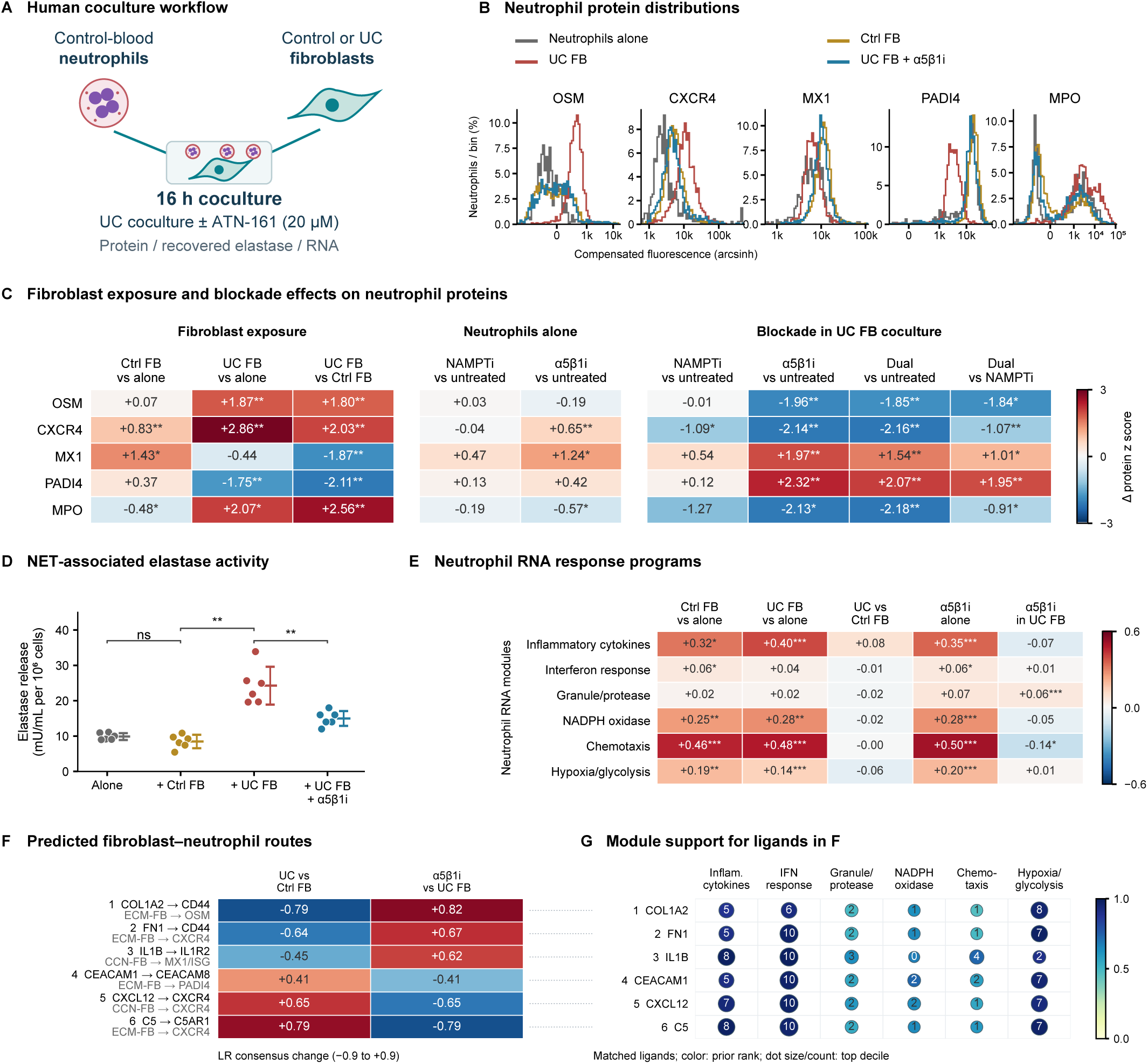
UC fibroblasts increase neutrophil inflammatory readouts that are attenuated by ATN-161. (A) Human coculture workflow: control-blood neutrophils with control or UC fibroblasts for 16 hours; UC cocultures with or without ATN-161 (20 µM), followed by protein, recovered elastase, and RNA measurements. (B) Representative compensated fluorescence histograms: gray, neutrophils alone; ochre, control fibroblasts; red, UC fibroblasts; blue, UC fibroblasts plus ATN-161. (C) Differences in sample-median neutrophil fluorescence, scaled by marker-specific SD across 47 samples; n=6 per arm except control fibroblast coculture (n=5). Exact unpaired permutation tests are BH-adjusted across 45 comparisons. MX1/PADI4 channel identities remain provisional. (D) NET-associated elastase activity; six independent experiments per arm, mean±SD. Three exact unpaired comparisons use Holm correction. All arms use the same scale and normalization. (E) Six RNA programs; mean paired differences across five contrasts (n=5,6,5,5,5 recorded donors; ≥20 neutrophils per arm). Paired t tests are BH-adjusted across 30 tests; exact sign-flip sensitivity tests yielded no BH discoveries. (F) Six predicted routes selected for opposite exposure/blockade directions. (G) Prior-based ligand–module support: color, mean gene percentile; size and labels, top-decile target counts. These predictions do not establish receptor-specific signaling. ATN-161 (α5β1i), α5β1-directed inhibitor; FK-866 (NAMPTi), NAMPT inhibitor; dual, both; FB, fibroblasts; Ctrl, control. *Adjusted P<0.05; **<0.01; ***<0.001; ns, ≥0.05.

We then assessed whether the protein changes were accompanied by an effector-associated response. Across six independent experiments per group on a common measurement scale, NET-associated elastase activity was higher with UC than control fibroblasts and lower after ATN-161 within UC coculture (both Holm-adjusted P = .00649; Figure 3D). Control fibroblasts versus neutrophils alone were nonsignificant (adjusted P = .132). Together, these experiments show that UC fibroblast exposure increases neutrophil inflammatory proteins and NET-associated elastase activity, and that these responses are pharmacologically modifiable.

### Coculture transcriptomics and communication analyses nominate candidate responses

Having established changes in protein and elastase measurements, we examined accompanying neutrophil RNA programs (Figure 3E) from coculture scRNA-seq. Fibroblast annotation and target-expression controls are shown in Supplementary Figure 4; neutrophil annotation and quality control are shown in Supplementary Figure 5. Paired parametric testing identified higher chemotaxis scores with control and UC fibroblasts than with neutrophils alone (+0.460 and +0.482 standardized units; q = .0000963 and .000308), together with inflammatory-cytokine, oxidase-associated, and hypoxia/glycolysis increases. Interferon increased with control fibroblasts (q = .0132), but the UC comparison was nonsignificant. No program distinguished UC from control fibroblasts after correction.

Within UC coculture, ATN-161 reduced the chemotaxis score by 0.137 standardized units (q = .0498) and increased the granule-associated score by 0.0627 units (q = .0000220); the other four programs changed nonsignificantly. ATN-161 alone increased several programs. Separate recorded-donor state-composition contrasts are shown in Supplementary Figure 6. Communication analyses nominated chemotactic, adhesion-associated, and cytokine routes selected for opposite directions with UC exposure and ATN-161 (Figure 3F). These included CXCL12–CXCR4, C5–C5AR1, CEACAM1–CEACAM8, COL1A2–CD44, FN1–CD44, and IL1B–IL1R2; IL1R2 is a decoy receptor.^29^ Ligand–module mapping linked these candidates to overlapping inflammatory, interferon, and metabolic target sets (Figure 3G). Communication analyses suggested selective remodeling of predicted fibroblast–neutrophil interactions with α5β1-directed treatment, involving chemotactic, adhesion-associated, and cytokine routes. These predictions provide candidate mechanisms for the observed protein and elastase responses, while prior-based mapping identifies potential downstream inflammatory, interferon, and metabolic programs for mechanistic investigation.

### FAP+ stromal ablation reduces colitis and recovered neutrophil representation

The coculture experiments established modifiable neutrophil responses to fibroblast exposure. We next examined how removing FAP+ cells affected tissue inflammation and neutrophil representation in DSS colitis. Ablation was associated with lower histologic injury and colonic neutrophil and fibroblast frequencies (Figure 4A). Milo analysis identified depleted Osm-, Padi4-, Mx1-, and Cxcr4-associated neighborhoods in 4 of 7, 5 of 6, 2 of 2, and 12 of 13 tested neighborhoods, respectively, with none enriched (Figure 4B and C). Il1r1-positive fibroblast neighborhoods were depleted in 16 of 16, while 9 of 15 Ackr4-positive Pi16-like neighborhoods were enriched. Absorptive– secretory transitional epithelium was enriched and IFN/MHC-II-responsive epithelium predominantly depleted. These are counts of overlapping neighborhoods, not independent state-level tests.

**Figure 4.**
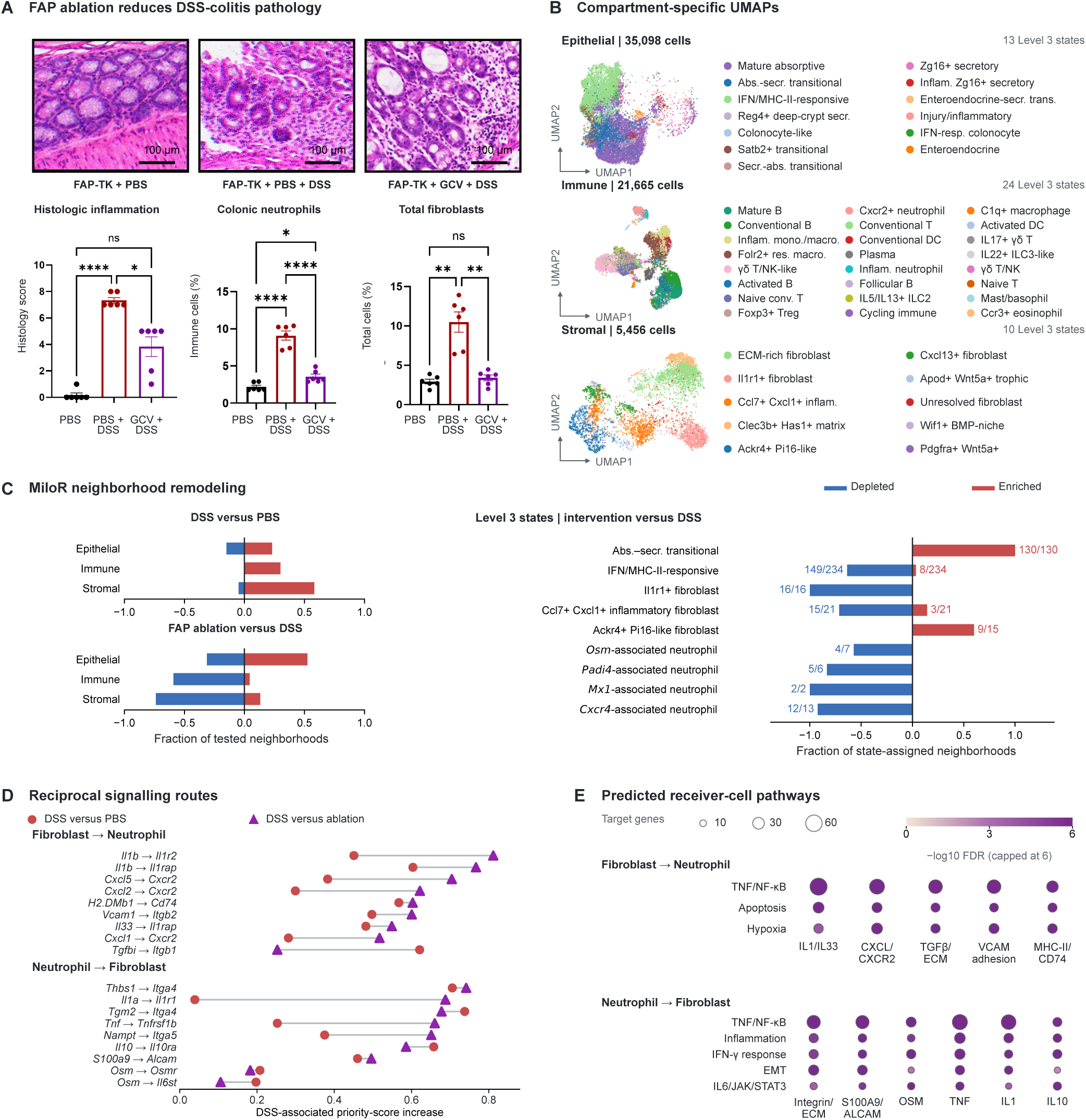
FAP+ stromal ablation attenuates colitis and changes cellular composition. (A) Histology, colonic neutrophil, and fibroblast measurements for FAP-TK+PBS, FAP-TK+PBS+DSS, and FAP-TK+ganciclovir+DSS. Histology was scored by a blinded pathologist. Displayed observations and assay-specific significance annotations are retained. Middle-field luminance sharpening is described in Supplementary Methods; 100-μm scale-bar calibration and field identities remain provisional. (B) Pooled UMAPs of 35,098 epithelial, 21,665 immune, and 5,456 stromal cells. (C) Fractions of Milo neighborhoods enriched or depleted at FDR<0.05; labels indicate significant/tested overlapping neighborhoods. Red, enrichment; blue, depletion. RNA-state labels do not denote protein-positive gates; pseudo-derived units remain in this model. (D) Predicted reciprocal-route differences: red circles, DSS-minus-control; purple triangles, DSS-minus-ablation. (E) Predicted receiver-pathway support; size indicates target overlap and color −log₁₀ BH FDR, capped at 6. Treatment–batch confounding and sparse residual neutrophils limit transcriptomic interpretation.

Flow cytometry showed lower OSM-, PADI4-, and MX1-positive percentages within neutrophils after ablation; the CXCR4 comparison was nonsignificant (Supplementary Figure 7C–F). Only two neutrophils were recovered from five ablation mice versus 1,074 from six DSS mice in the trajectory dataset, precluding within-state trajectory inference (Supplementary Figure 7A,B). These readouts support reduced neutrophil representation, with recruitment, persistence, survival, and cell recovery remaining possible contributors. Exploratory communication priorities were lower for fibroblast Cxcl–Cxcr2 recruitment inputs and reciprocal neutrophil Osm–Osmr/Il6st routes, with predicted targets overlapping inflammatory and stress-response programs (Figure 4D and E). Sparse residual neutrophils limit interpretation of these molecular predictions.

### α5β1-directed blockade lowers subset MPO despite higher marker-positive percentages

Because FAP ablation removes multiple stromal inputs, we next examined α5β1-directed pharmacologic treatment in a separate DSS experiment using ATN-161, an α5β1 inhibitor. ATN-161 was associated with lower histologic injury and overall colonic neutrophil and fibroblast frequencies (Figure 5A). MPO fluorescence was lower within OSM-positive and PADI4-positive neutrophils (Figure 5B), although the corresponding marker-positive percentages within neutrophils increased (Supplementary Figure 10C,D). CXCR4-positive and MX1-positive percentages and MPO within CXCR4-positive neutrophils changed nonsignificantly (Supplementary Figure 10E–G). Thus, lower overall neutrophil frequency and subset MPO coexisted with higher selected marker-positive percentages, without establishing absolute subset expansion. Detectable α5β1-positive fibroblast fractions also decreased, although receptor occupancy or epitope interference could contribute (Supplementary Figure 10A,B). Additional human protein-positive measurements are reported separately (Supplementary Figure 9).

**Figure 5.**
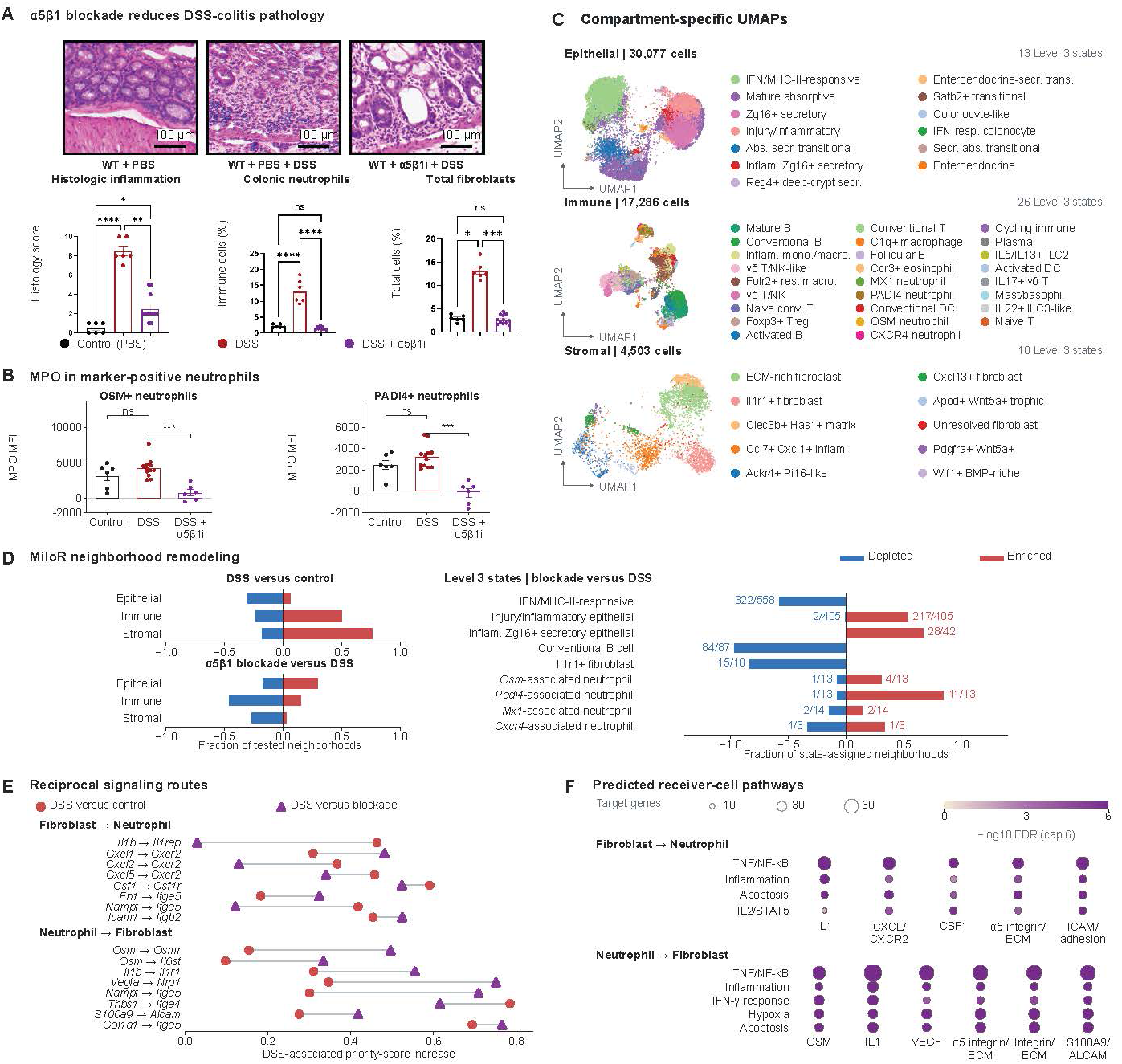
α5β1-directed treatment reduces colitis and neutrophil subset MPO. (A) Histology and colonic neutrophil and fibroblast frequencies in control, DSS+PBS and DSS+ATN-161 mice. Histology was scored blinded. Image processing, provisional field identities and 100-μm scale calibration are detailed in Supplementary Methods. (B) MPO fluorescence within OSM-positive and PADI4-positive neutrophils. A,B retain source observations, error bars and annotations; control-versus-DSS MPO comparisons use reconstructed-point Welch/BH tests, with other comparisons retaining source tests. (C) UMAPs of 30,077 epithelial, 17,286 immune and 4,503 stromal cells; RNA inventory, 6 control, 6 DSS and 12 blockade units. (D) Fractions of Milo neighborhoods depleted/enriched at FDR<0.05; labels, significant/tested overlapping neighborhoods. (E) Predicted route differences: red circles, DSS-minus-control; purple triangles, DSS-minus-blockade. Three blockade mice meet communication eligibility. (F) Predicted receiver-pathway support; size, target overlap; color, −log₁₀ BH FDR, capped at 6. One combination has FDR=0.0622. Supplementary Figures 8 and 10 provide trajectories, marker-positive percentages and CXCR4-subset MPO. Treatment–batch confounding and assay-specific statistical provenance are described in Supplementary Methods.

To place these tissue and protein responses in a cellular context, we next examined compartment-specific RNA profiles (Figure 5C) and neighborhood representation (Figure 5D). The blockade-associated pattern included both enrichment and depletion: Osm-associated neighborhoods comprised 4 of 13 enriched and 1 of 13 depleted neighborhoods, and Padi4 comprised 11 of 13 enriched and 1 of 13 depleted. Mx1 and Cxcr4 categories also contained both directions. Il1r1-positive fibroblast and IFN/MHC-II-responsive epithelial neighborhoods were predominantly depleted, while injury-associated and Zg16-positive secretory epithelial neighborhoods were enriched.

Cross-sectional trajectory analysis provided a complementary description of recovered neutrophils (Supplementary Figure 8). Among 2,147 cells, α5β1 blockade samples occupied the Cxcr2-associated region more frequently than DSS samples (approximately 94% versus 43%). Transferred CXCR4-associated state fractions decreased from 29.7% to 10.3% (−19.4 percentage points; q = .0196); OSM/PADI4 increases and MX1 changes were nonsignificant. These occupancy differences do not demonstrate reversal of individual cells. Transferred RNA programs, Milo neighborhoods, and protein-positive gates describe distinct classifications.

Communication predictions again nominated Cxcl–Cxcr2 recruitment, adhesion-associated, and reciprocal Osm/Il1 cytokine routes, with fibroblast target sets including Ccl2 and Cxcl5 (Figure 5E and F). These candidates provide molecular context for the intervention-associated patterns.

### Stromal interventions share epithelial inflammatory changes but differ in differentiation responses

The reductions in colitis severity and neutrophil frequency prompted us to examine accompanying epithelial responses. Relative to the DSS-only reference, whole-epithelium inflammatory chemokine scores were lower after FAP ablation and α5β1 blockade (q = .0229 and .000431). Ablation was associated with higher absorptive differentiation and lower injury-associated regeneration scores (q = .0327 and .0229), whereas blockade was associated with higher barrier/junction and mucus/secretory scores and lower absorptive differentiation (q = .0269, .000431 and .000431; Figure 6A). The ablation-associated mucus increase did not meet the adjusted significance threshold (q = .100). These comparisons describe intervention-associated patterns from separate experiments.

**Figure 6.**
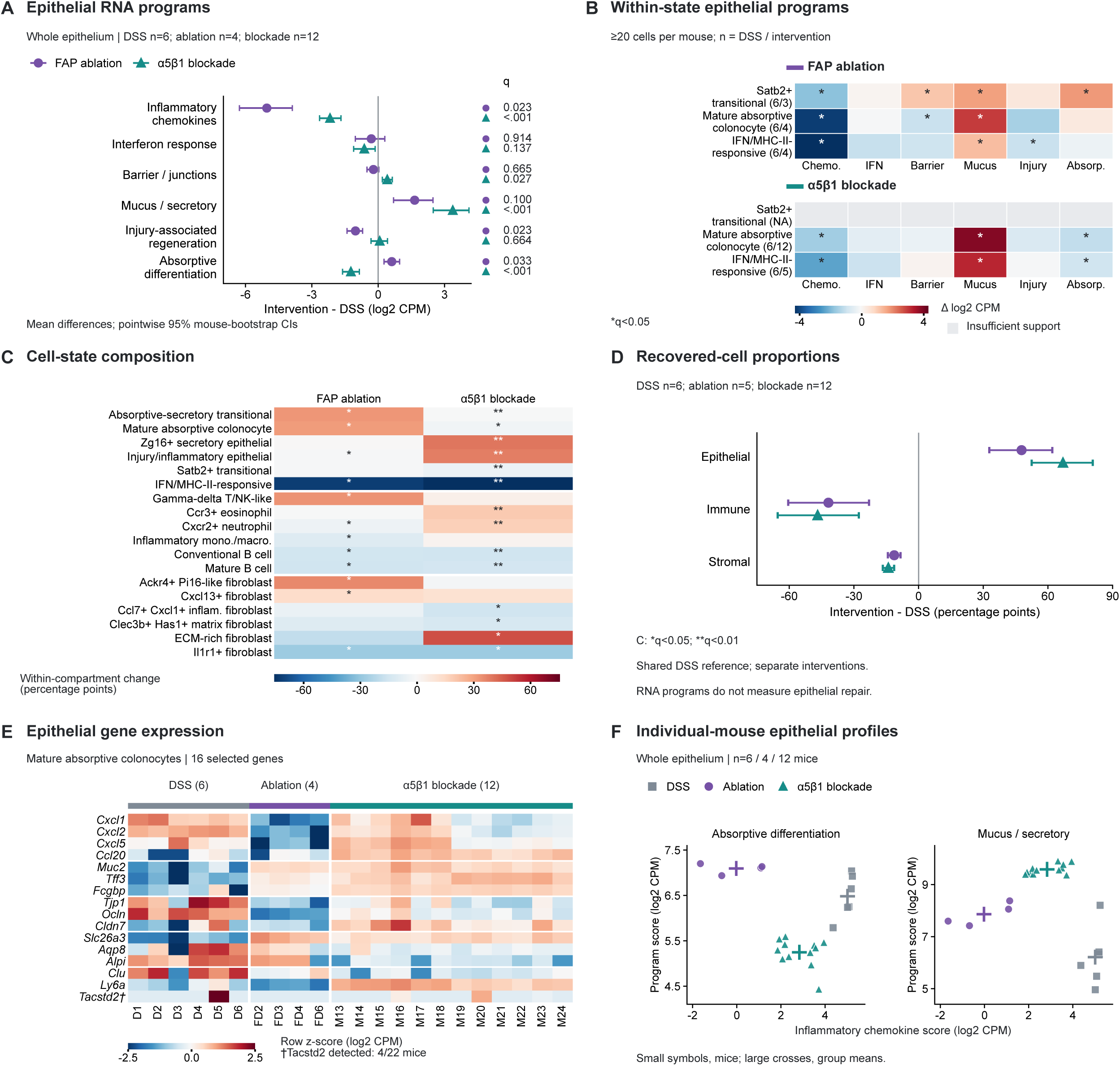
Shared and distinct epithelial responses to stromal interventions. (A) Whole-epithelium RNA-program differences versus DSS. Points and lines show mean differences and pointwise 95% mouse-bootstrap confidence intervals. (B) Within-state program differences; labels give DSS/intervention mouse numbers. States require ≥20 cells/mouse and ≥3 mice/group; gray/NA indicates insufficient support. (C) Within-compartment proportion changes for six previously selected states per compartment; white separators distinguish epithelial, immune and stromal states. (D) Recovered-cell compartment-proportion differences with pointwise 95% confidence intervals from 10,000 mouse bootstraps. Purple circles, FAP ablation; teal triangles, α5β1 blockade. (E) Mature absorptive-colonocyte expression of 16 selected genes. Columns represent individual mice; colors show gene-wise z-scores of stored log₂ CPM, saturated at ±2.5. †Tacstd2 was detected in 4/22 mice. (F) Existing whole-epithelium chemokine scores versus absorptive and mucus/secretory scores. Small symbols represent mice; crosses, group means; gray squares, DSS. For A,E,F, DSS/ablation/blockade n=6/4/12 biological mice; for C,D, n=6/5/12. Mann-Whitney tests with BH correction across 12 comparisons (A), 30 supported comparisons (B), or all states per intervention (C); *q<0.05, **q<0.01. Chemo., inflammatory chemokines; IFN, interferon; Injury, injury-associated regeneration; Absorp., absorptive differentiation. Panels E,F are descriptive, without additional hypothesis tests. Separate intervention experiments share a DSS reference; treatment and acquisition batch are confounded. Proportions do not establish absolute abundance, and RNA programs do not establish epithelial function or repair. Only biological mice are included in the displayed comparisons.

To assess whether these patterns reflected changes beyond broad epithelial-state proportions, we examined programs within supported states. Both interventions were associated with lower chemokine and higher mucus/secretory scores in mature absorptive colonocytes and IFN/MHC-II-responsive epithelial cells (all corresponding q < .05; Figure 6B). Junction responses were state dependent: the mature-colonocyte barrier/junction score was lower after ablation (q = .0238), while the blockade contrast did not meet the adjusted threshold (q = .0655). Thus, the whole-epithelium barrier increase under blockade did not represent uniform junction-program elevation across states.

The mature-colonocyte gene display showed selective expression patterns across chemokine, mucus, junction and differentiation genes, including higher mucus-associated expression and lower *Alpi* expression in the blockade group (Figure 6E). The complete 39-gene display retains the original program membership (Supplementary Figure 11). Individual-mouse profiles showed lower chemokine scores with higher absorptive scores in the ablation group, whereas blockade combined lower chemokine scores than DSS with higher mucus/secretory and lower absorptive scores (Figure 6F). These descriptive displays expose individual variation and gene-level heterogeneity without additional significance tests or evidence of functional restoration.

Cellular composition provided complementary context. Both interventions were associated with higher epithelial and lower immune and stromal fractions among recovered cells (Figure 6D). Ablation increased absorptive-secretory transitional and mature absorptive representation, whereas blockade increased Zg16-positive secretory and injury/inflammatory epithelial representation; IFN/MHC-II-responsive representation was lower with both (Figure 6C). Across 47 shared states, 29 changed in the same direction, but concordance was modest and nonsignificant (Spearman ρ = 0.285; P = .052; Supplementary Figure 12A). These proportions do not establish absolute epithelial expansion. Exploratory communication priorities were commonly lower despite the different cellular and epithelial patterns: of 691 matched routes, 502 were jointly DSS-induced and 490 of these received lower priority after both interventions (Supplementary Figure 12B and C). These predicted changes provide candidate molecular context rather than measured signaling activity.

### Independent human data identify activated fibroblasts with relevant receptor components

Finally, we asked whether independent human datasets supported the activated stromal features implicated by the tissue and experimental findings. We examined TAURUS, a longitudinal single-cell atlas of intestinal biopsies from patients with UC and Crohn’s disease sampled before and after treatment with the anti-TNF antibody adalimumab.^26^ In the UC subset, inflammation was associated with increases in activated FAP-positive fibroblast fraction, FAP-inflammatory programs, and α5β1-adhesion programs (q = .023, .039, and .023, respectively; Figure 7A). Activated fibroblasts had higher adhesion scores than other fibroblasts in within-patient comparisons across 16 patients (Figure 7B). Across 50 eligible biopsies, ITGA5, ITGB1, OSMR, and IL6ST expression was higher in activated fibroblasts (all q < .005). Same-cell ITGA5/ITGB1 and OSMR/IL6ST transcript codetection increased by 14.57 and 14.61 percentage points, respectively (both q = .000641; Figure 7C), supporting enrichment of adhesion- and cytokine-receptor components.

**Figure 7.**
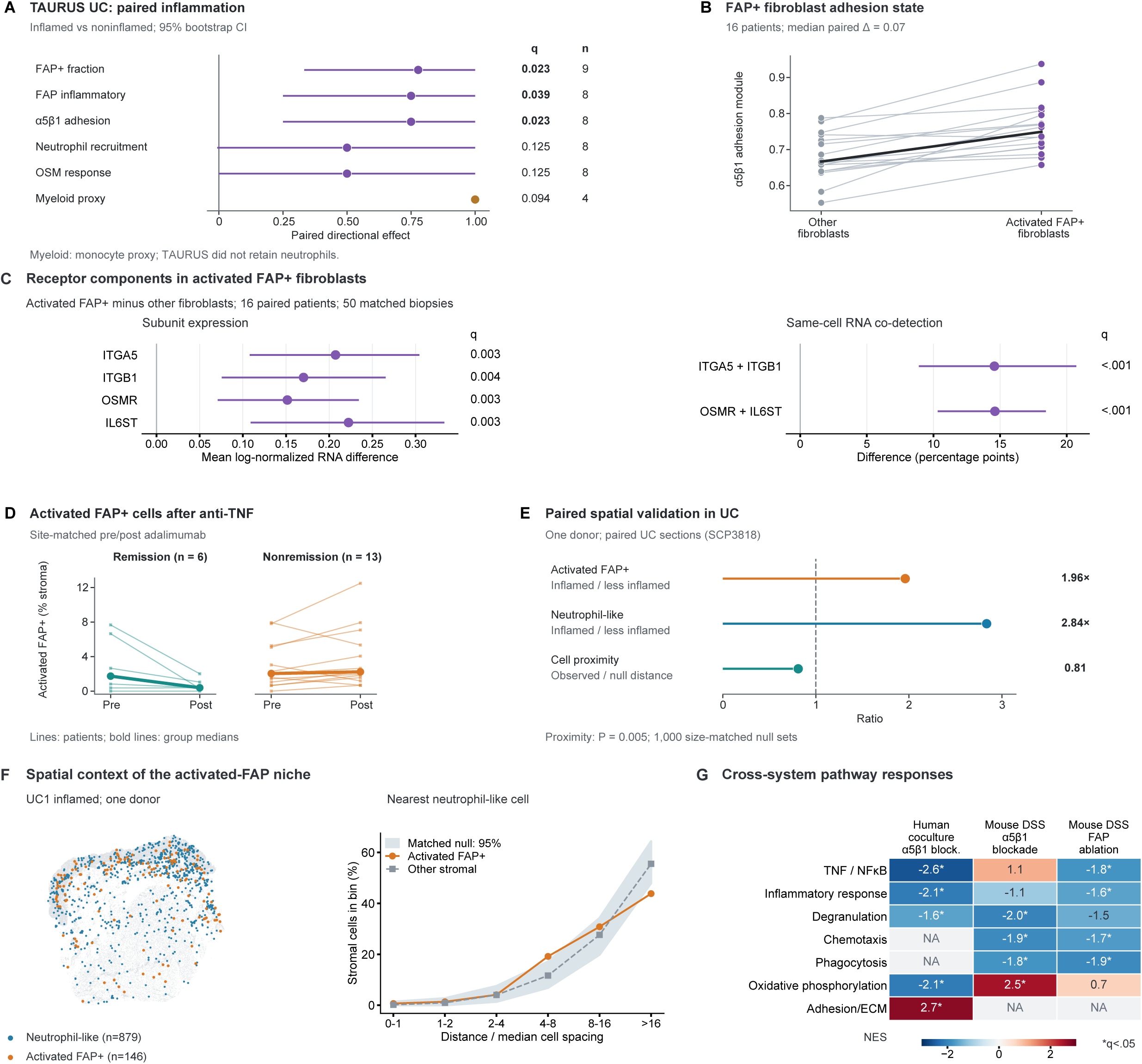
Independent human data support activated fibroblast receptor-component enrichment. (A) TAURUS paired inflammation-associated sign effects and 95% bootstrap CIs; feature-specific n=9,8,8,8,8,4 patients; BH correction across six features. The myeloid feature is a monocyte proxy. (B) Activated-versus-other fibroblast adhesion scores in 16 patients; thin lines, pairs; bold line, medians. (C) Patient-averaged receptor expression and transcript-codetection differences from 50 biopsies in 16 patients; ≥5 activated and ≥20 other fibroblasts per biopsy. Lines show 95% patient-bootstrap CIs; exact sign-flip tests are BH-adjusted across six outcomes. (D) Site-matched adalimumab trajectories; 6 remission and 13 nonremission patients; bold lines, medians. (E) Paired SCP3818 sections from one donor, with a size-matched stromal distance null. (F) Inflamed-section map and normalized distances for 146 activated-FAP and 879 neutrophil-like cells; null stromal sets match local epithelial-neighbor fraction. E,F use 1,000 randomizations and distinct nulls. (G) Cross-system normalized enrichment scores; NA, unavailable; stars, source q<0.05. RNA codetection does not establish assembled receptors; spatial observations represent one donor.

Site-matched adalimumab trajectories were heterogeneous and did not establish a response predictor (Figure 7D). Paired spatial sections from one UC donor showed higher activated-FAP and neutrophil-like fractions in inflamed tissue and closer proximity than size-matched random stromal sets (distance ratio, 0.811; empirical P = .004995; Figure 7E). An epithelial-neighborhood-matched null provided additional anatomical context (Figure 7F). Cross-system pathway summaries combined different populations and contrasts (Figure 7G). These findings corroborate selected human stromal observations. Absent TAURUS neutrophils and the single spatial donor preclude independent validation of the complete experimental mechanism.

## Discussion

This study identifies a pharmacologically modifiable influence of colonic fibroblast exposure on neutrophil inflammatory phenotype and effector-associated activity. UC fibroblasts increased neutrophil OSM, CXCR4 and MPO fluorescence and NET-associated elastase activity relative to control fibroblasts, and α5β1 blockade attenuated these responses. Human spatial profiling linked FAP-high fibroblast proximity to higher neutrophil OSM and CXCR4 fluorescence. In mice, stromal depletion and integrin-directed treatment reduced histologic inflammation and overall neutrophil frequency while producing different neutrophil protein and representation patterns. The accompanying epithelial analyses extend the tissue context: both interventions were associated with lower chemokine programs but distinct absorptive and secretory responses. Together, these findings support fibroblast-associated pathways as candidate targets for modifying mucosal inflammation in UC.

The coculture experiments are the principal experimental advance beyond the tissue associations. Prior studies established OSM-responsive intestinal stroma and IL-1-dependent fibroblast recruitment of neutrophils,^8,9^ and our published atlas identified neutrophil states and candidate stromal circuits associated with treatment resistance.^12^ Exposure of control-blood neutrophils to UC fibroblast preparations altered both inflammatory proteins and an effector-associated readout outside the tissue environment. These results extend the recruitment framework to modulation of neutrophil inflammatory properties. The concordant attenuation of protein and elastase measurements supports the therapeutic relevance of this response, while the cellular site and mechanism of drug action remain unresolved.

FAP depletion and integrin-directed treatment produced distinct patterns that are relevant to interpreting mucosal interventions. FAP+ cell removal changes multiple stromal inputs, whereas ATN-161 binds multiple integrins^30^ and can act on several cellular compartments. Distinct inflammatory and remodeling functions of fibroblast subsets in other tissues further support considering the nature of the stromal perturbation.^31^ After ATN-161, higher OSM-positive and PADI4-positive percentages within neutrophils coexisted with lower MPO fluorescence within those subsets and lower overall neutrophil frequency (Figure 5A,B; Supplementary Figure 10C,D). The subset MPO measurements therefore add a protein-phenotype readout to the reduction in tissue inflammation. Together, these findings support assessing cellular representation and inflammatory proteins separately when evaluating stromal-directed interventions. Neither marker-positive percentages nor MPO fluorescence alone establishes effector function.

Exploratory RNA and communication analyses nominate recruitment, matrix-associated, and reciprocal cytokine pathways for mechanistic investigation. Their evidential weight is limited by test-sensitive coculture RNA findings and the distinction between predicted ligand–target relationships and measured signaling. These analyses provide candidate explanations for the protein and elastase responses, with the strongest disease-associated coculture differences observed in those experimental readouts.

The spatial findings place this experimental response in a plausible human tissue context: FAP-high fibroblast proximity was associated with selected neutrophil proteins, and external human data supported activated-fibroblast receptor-component enrichment. The absence of corrected associations with operational α5β1-positive proximity limits a receptor-specific spatial interpretation. Although stromal apposition can influence epithelial phenotypes in gastric preneoplasia,^32^ proximity in UC does not resolve whether direct contact, soluble mediators, or matrix-associated inputs underlie the neutrophil observations.

The epithelial findings connect stromal intervention to the compartment responsible for mucosal absorption and barrier maintenance. Shared reductions in chemokine programs accompanied different differentiation-associated responses: ablation was associated with higher absorptive and lower injury-associated regeneration scores, whereas blockade was associated with higher mucus/secretory scores and lower absorptive scores (Figure 6A). Lower chemokine and higher mucus scores within defined epithelial states indicate that these patterns were not solely attributable to shifts in broad state proportions (Figure 6B). The gene and individual-mouse displays make the heterogeneity visible, including junction genes that did not increase uniformly (Figure 6E and F). They reinforce the interpretation of distinct epithelial responses rather than establish direct fibroblast-to-epithelial signaling or a neutrophil-mediated causal pathway.

Therapeutic development must therefore distinguish control of inflammation from preservation of epithelial function and host defense. Higher mucus or junction RNA scores do not establish improved secretion, permeability or repair, and the different absorptive patterns warrant functional evaluation. Neutrophil-derived IL-22 can support epithelial restitution,^33^ PAD4-dependent immunothrombi can limit mucosal bleeding,^34^ and stromal MAP3K2–R-spondin1 signaling contributes to regeneration after intestinal injury.^35^ Alongside studies of stromal adhesion targets in Crohn’s disease,^7^ these observations support assessing epithelial differentiation and tissue-supporting consequences together with neutrophil abundance, inflammatory proteins and effector-associated activity.

The principal limitations concern attribution and the scope of the mechanistic readouts. ATN-161 pharmacology does not establish a fibroblast-specific α5β1 requirement. Although elastase and MPO participate in NET formation,^36^ elastase activity alone does not establish NET structure, and migration, adhesion, antimicrobial function and the mechanism of neutrophil loss were not directly measured. Epithelial RNA programs are investigator-curated expression summaries. External human datasets corroborate selected stromal findings rather than the complete mechanism. Acute DSS colitis does not establish efficacy in chronic UC or reversal of treatment resistance.

By linking human tissue observations with coculture perturbations and complementary DSS colitis experiments, this study extends fibroblast regulation of neutrophils to inflammatory phenotype and effector-associated activity. The coculture experiments demonstrate modifiable responses to fibroblast exposure, while the mouse interventions connect reduced histologic inflammation to distinct neutrophil and epithelial patterns. Their shared epithelial chemokine reduction and divergent differentiation-associated programs suggest that control of inflammation can accompany different tissue responses. Together, these findings identify fibroblast-associated pathways as candidate therapeutic targets for UC and support evaluating epithelial programs alongside neutrophil frequency, subset proteins and functional measurements. This approach could guide tissue pharmacodynamic assessment of stromal-directed therapies. Translation will require identifying the responding cellular targets and establishing whether inflammation can be reduced while preserving host defense, epithelial function and tissue repair.

## Data and code availability

The published discovery atlas is available through the Broad Single Cell Portal (SCP3755). TAURUS data are available in Zenodo (version 3; DOI: 10.5281/zenodo.14007626), and the independent spatial dataset is available through SCP3818. Analysis scripts, selected derived source tables, and main and supplementary figures are publicly available in the Gubatan Lab GitHub repository (https://github.com/GubatanLab/UC-fibroblast-neutrophil-manuscript), with release 1.0.0 archived in Zenodo (DOI: <u>10.5281/zenodo.22881923</u>). The underlying CODEX datasets and human fibroblast–neutrophil coculture and mouse colitis scRNA-seq datasets are available from the corresponding author upon request. Dataset accessions, analysis-specific eligibility criteria, and the scope of deposited materials are detailed in the Supplementary Methods.

## Author contributions

Y.Z. and J.C.: Bioinformatic analyses and manuscript drafting, review, and revision. J.Y.: Investigation, methodology and validation for blood flow-cytometric analyses performed in the laboratory of S.S.; S.S.: Supervision of the blood flow-cytometric analyses. S.F.-S., D.R.H., and G.V.: Investigation and methodology for CODEX spatial multiplexed protein profiling. D.R.H.: Data curation, investigation and CODEX spatial multiplexed protein profiling, and writing—review and editing. G.N.: Supervision, resources, investigation and methodology for CODEX spatial multiplexed protein profiling. R.S., J.N.H. and T.F.: Participant recruitment, sample processing and single-cell experiments. Y.H. and K.P.: Results analyses and interpretation; manuscript drafting, review, and revision. S.R.: Conceptualization, funding acquisition, investigation, project administration, resources, supervision, validation, writing—original draft, and writing—review and editing; and participant recruitment, sample processing and single-cell experiments. J.G.: Conceptualization, data curation, analyses, investigation, methodology, supervision, validation, visualization and figure preparation, leadership of manuscript drafting, writing—original draft, and writing—review and editing.

## Declaration of AI-assisted manuscript preparation

Generative AI was not used in manuscript preparation.

## Supplementary figure legends

**Supplementary Figure 1.**
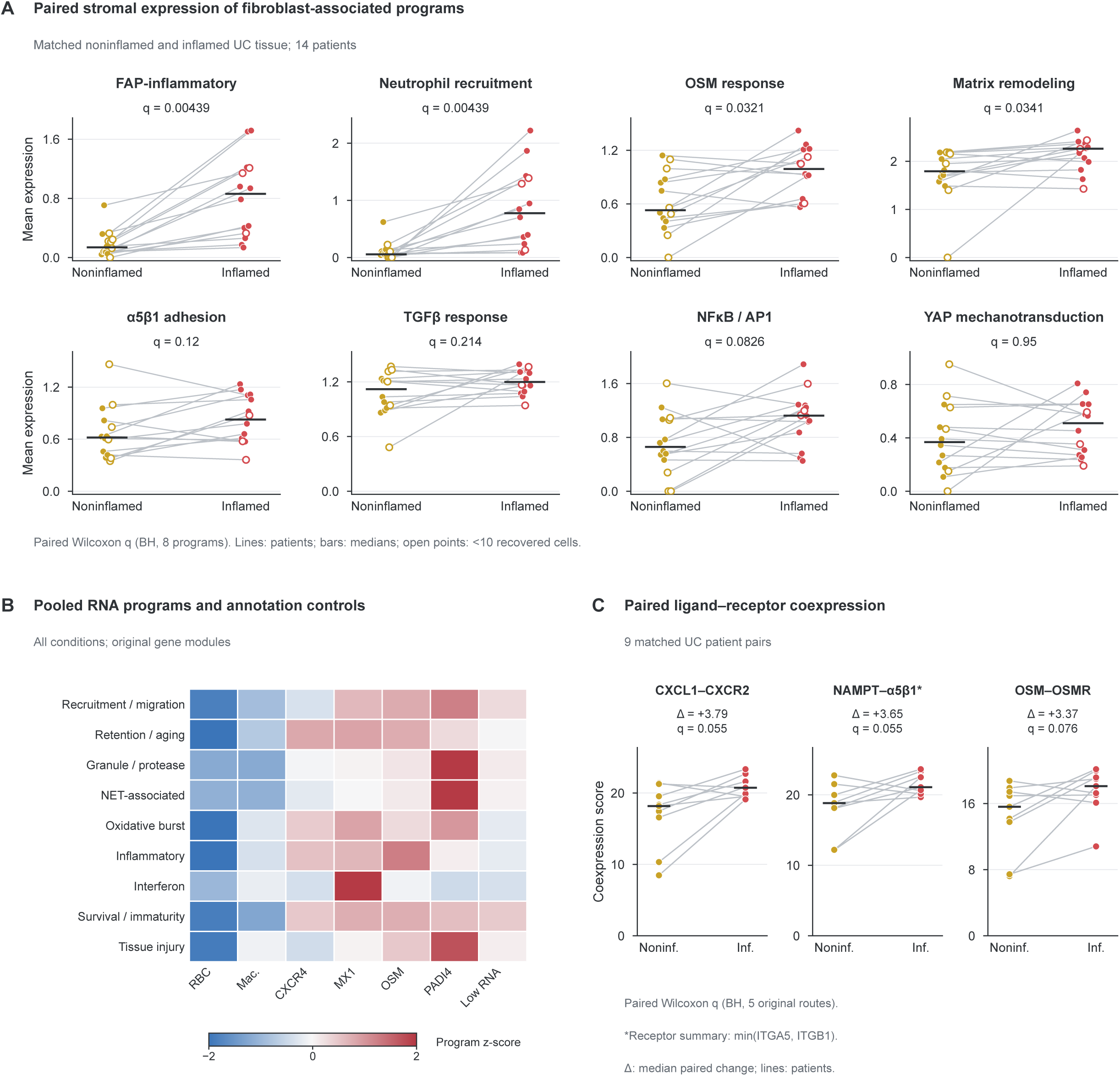
Stromal programs, neutrophil annotation controls, and paired coexpression. (A) Eight fibroblast-associated programs in the broader stromal compartment, including smooth-muscle cells and pericytes; 14 paired UC patients. Lines connect samples, bars indicate medians, and open points mark <10 cells. Paired Wilcoxon tests are BH-adjusted across eight programs. (B) Pooled original program expression, z-scored across seven annotations within each program. Counts: erythrocytes, 2; macrophages, 7; CXCR4, 9,614; MX1, 5,222; OSM, 8,432; PADI4, 5,911; low RNA, 9,505. The non-neutrophil annotations are controls. This cell-weighted display retains naming genes and differs from Figure 1E. (C) Coexpression in nine patient pairs; paired Wilcoxon tests retain BH correction across five original routes. Displayed q values are 0.055, 0.055, and 0.076; none is significant at q<0.05. The NAMPT receptor summary uses the lower ITGA5/ITGB1 expression and does not establish receptor assembly or binding.

**Supplementary Figure 2.**
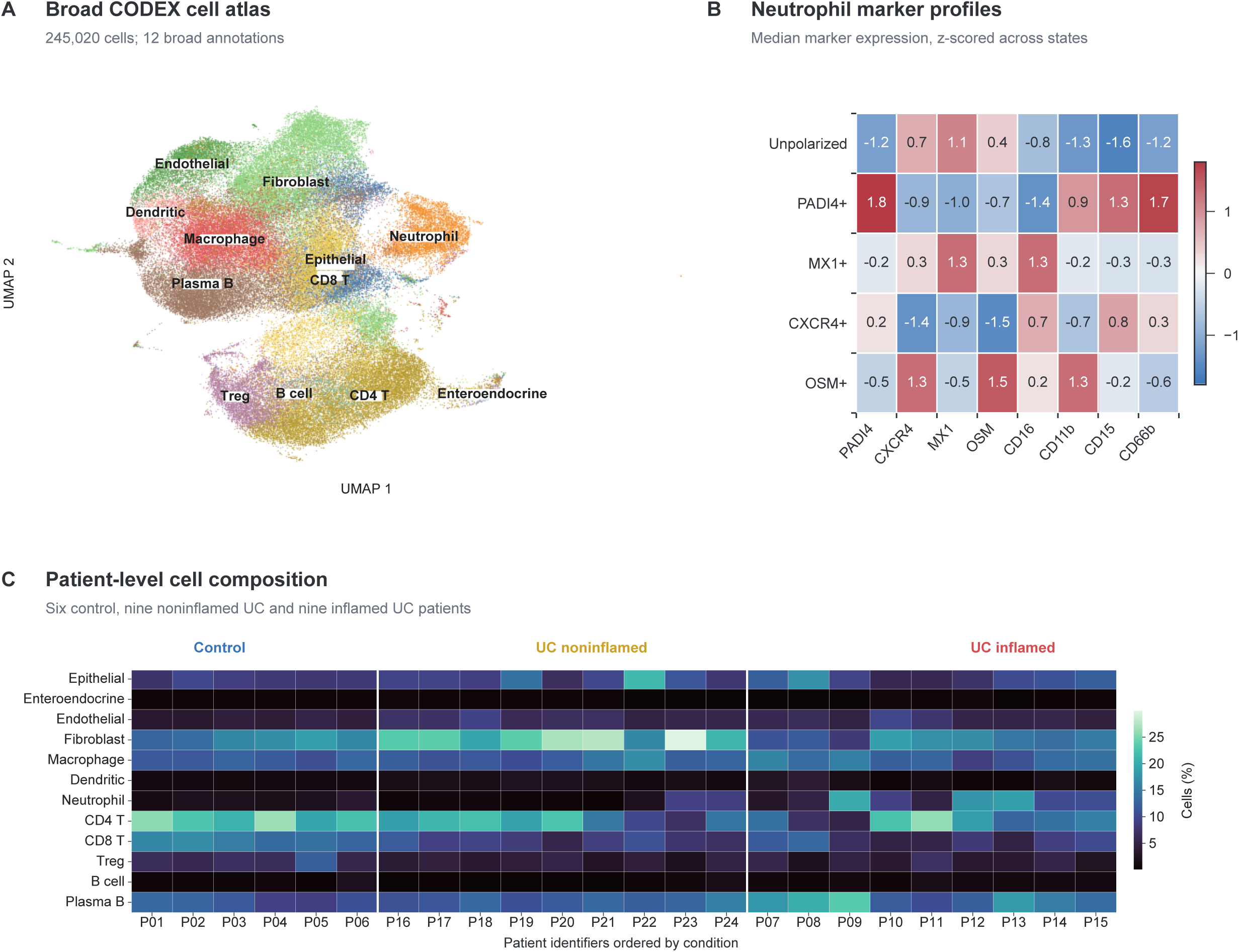
CODEX annotations and patient composition. (A) UMAP of 245,020 cells across 12 broad annotations. (B) Eight-marker median fluorescence across five neutrophil phenotypes, z-scored within marker; 14,906 neutrophils. (C) Cell-type fractions across 24 independent patients: 6 control, 9 noninflamed UC, and 9 inflamed UC. These descriptive summaries support Figure 2; cells are not independent replicates.

**Supplementary Figure 3.**
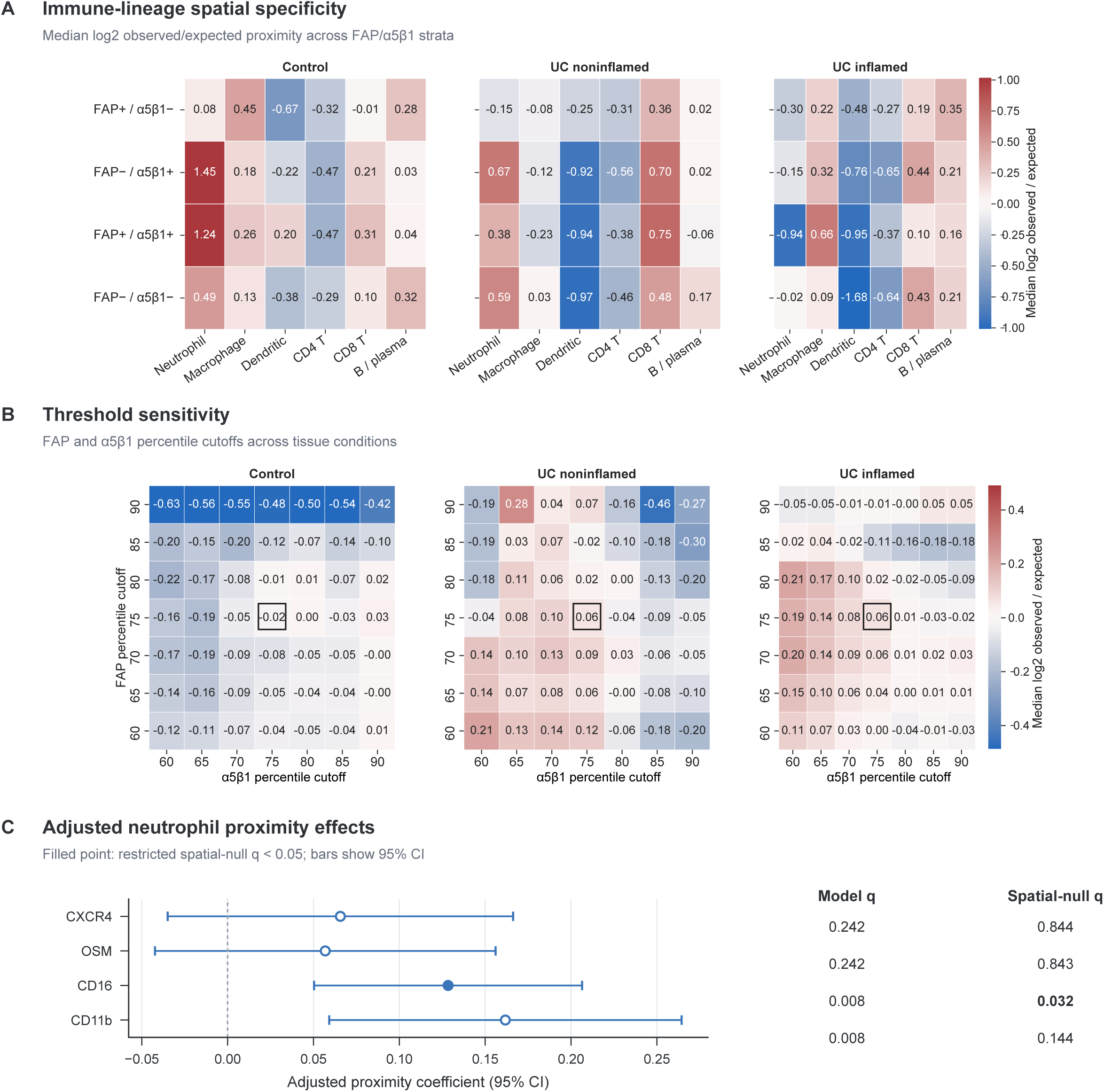
Spatial specificity and marker-threshold sensitivity. (A) Patient-median log₂ observed/expected enrichment of six immune lineages within 50 μm of four fibroblast categories; nulls retain coordinates and target abundance. (B) FAP-positive/α5β1-low proximity across 49 threshold combinations; outlines mark the 75th/75th percentiles. A,B include 6 control, 9 noninflamed, and 9 inflamed UC patients. (C) Adjusted coefficients and pointwise 95% CIs from 17 UC patients with ≥30 neutrophils; BH q values for covariate-adjusted models and 499 restricted permutations. Only CD16 passes both corrections. These reference definitions differ from Figure 2E.

**Supplementary Figure 4.**
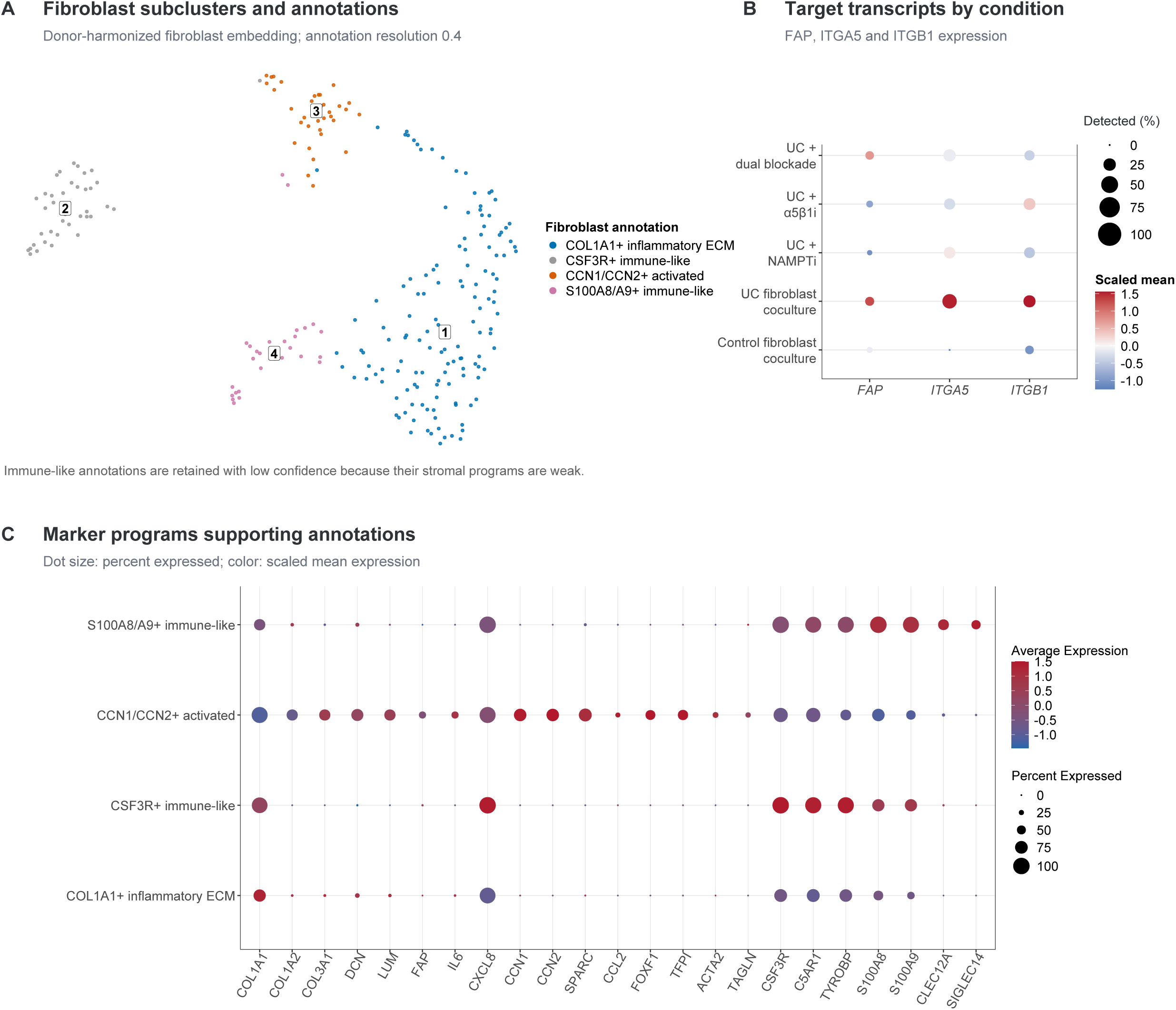
Fibroblast annotation and coculture transcript controls. (A) Fibroblast embedding after DecontX, principal-component analysis, Harmony, and Leiden clustering; 238 audit cells, including low-confidence immune-like controls. Communication analyses use 170 bona fide fibroblasts. (B) FAP, ITGA5, and ITGB1 across five fibroblast-containing conditions; size, detection; color, scaled expression. (C) Marker-program support for four annotations. Values use corrected log-normalized RNA. These descriptive controls have no hypothesis tests; sparse recovery limits condition-specific inference, and transcript codetection does not establish surface integrins.

**Supplementary Figure 5.**
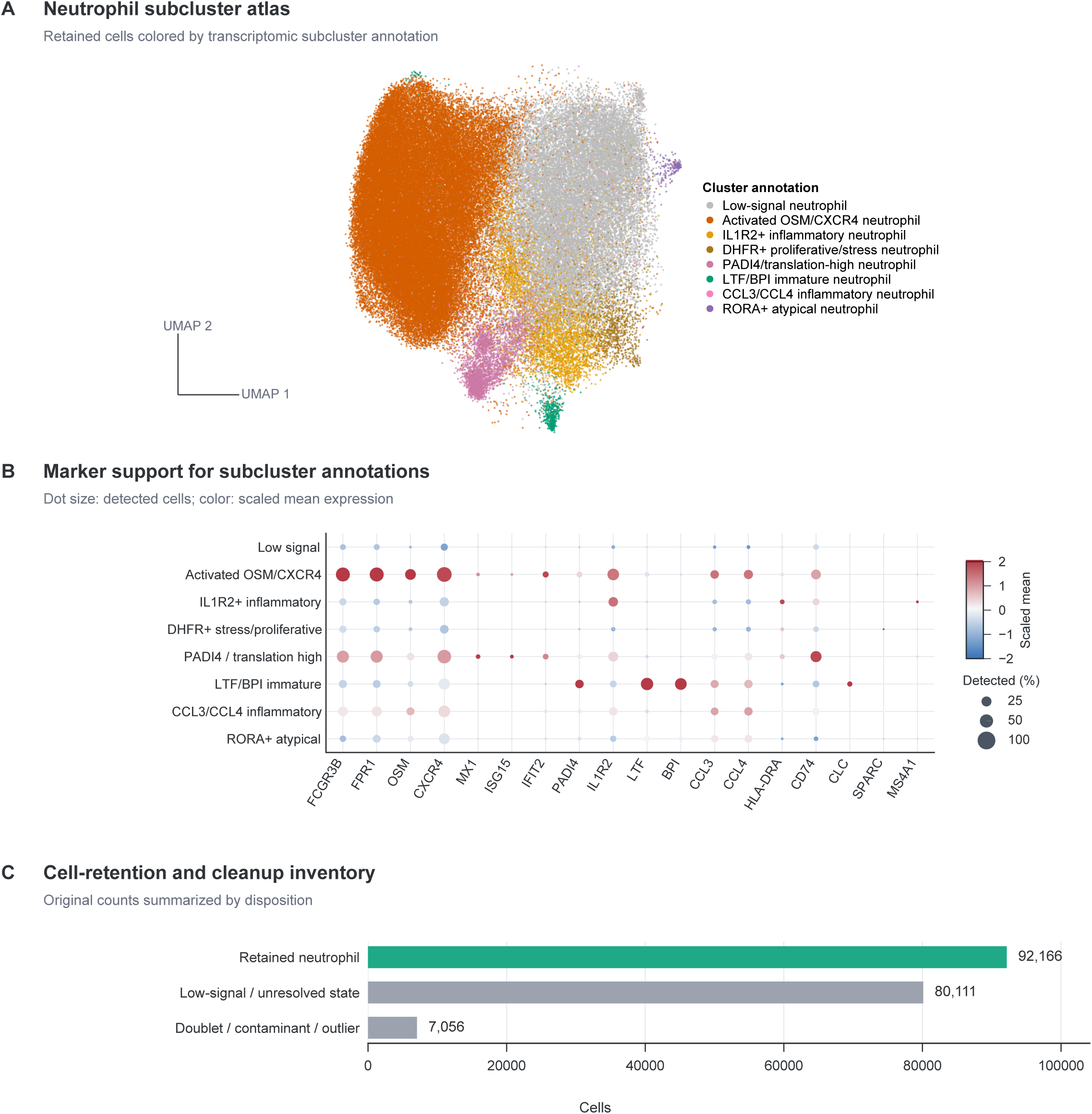
Neutrophil annotation and quality control. (A) Neutrophil-only scVI–Harmony embedding, including the low-signal audit group. (B) Eight subcluster annotations; size, marker detection; color, scaled expression (−2 to +2). (C) Cleanup inventory: 92,166 resolved neutrophils, 80,111 low-signal/unresolved cells excluded from state analyses, and 7,056 contaminant-, doublet-, or outlier-like cells removed. Counts describe different annotation stages and differ from protein-assay denominators.

**Supplementary Figure 6.**
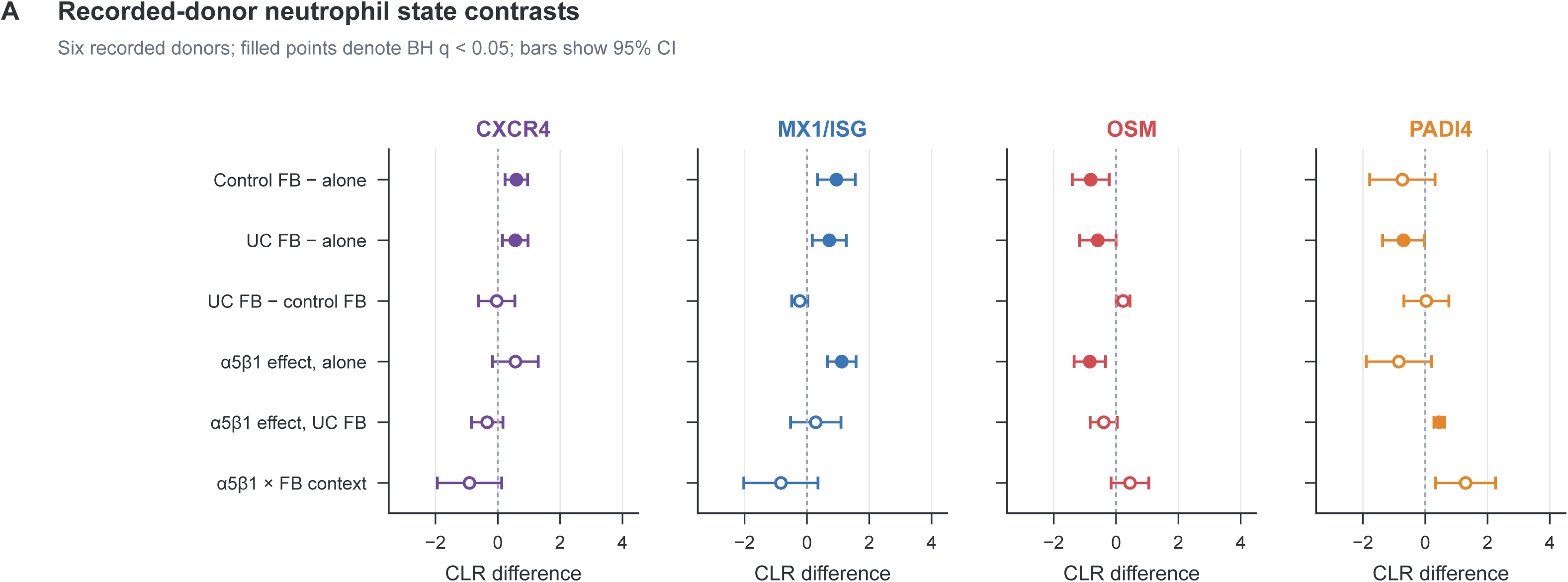
Recorded-donor neutrophil state contrasts. (A) Four-state centered-log-ratio effects across six contrasts; means and 95% parametric CIs using donor-specific variance. Six recorded donors contribute per comparison; no positive minimum-cell filter was applied. Filled/open symbols indicate BH q<0.05/≥0.05. The α5β1-by-context interaction is the inhibitor effect with UC fibroblasts minus its effect without fibroblasts; none of four state interactions passes correction. Biological matching remains unverified. These composition analyses are separate from Figure 3 protein and RNA-program tests.

**Supplementary Figure 7.**
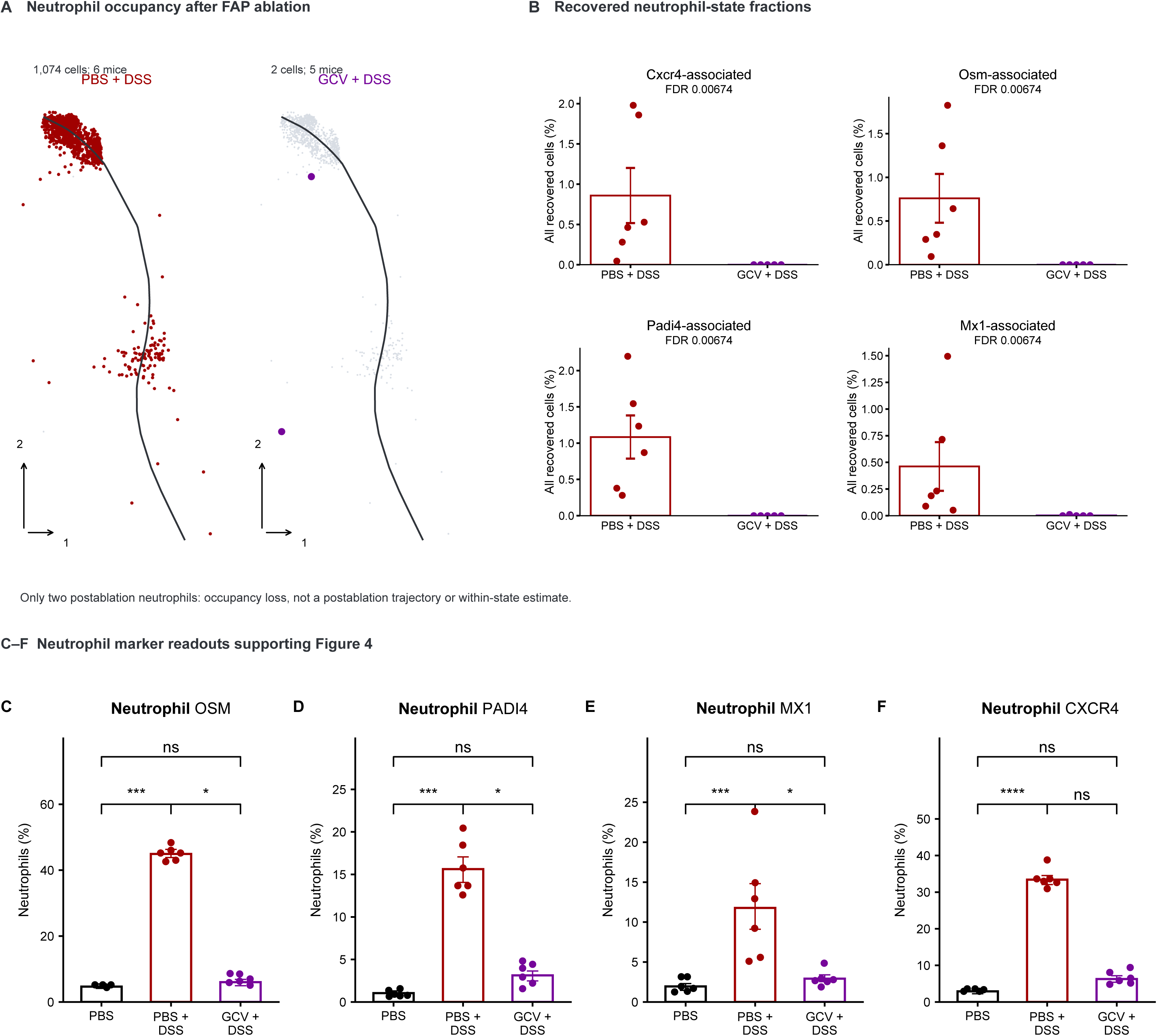
Neutrophil recovery and protein markers after FAP+ stromal ablation. (A) Shared manifold occupancy: DSS, 1,074 cells from six mice; ablation, two cells from five mice. Gray denotes the reference; curves do not measure time. (B) State fractions among all recovered cells per mouse; mean±SEM; Wilcoxon rank-sum tests with BH correction across four states. (C–F) OSM-, PADI4-, MX1-, and CXCR4-positive neutrophil measurements; black, PBS; dark red, PBS+DSS; purple, ganciclovir+DSS. Source observations, summaries, and significance annotations are retained. Sparse residual cells preclude within-state inference. Treatment–batch confounding and assay-specific statistical provenance are described in Supplementary Methods.

**Supplementary Figure 8.**
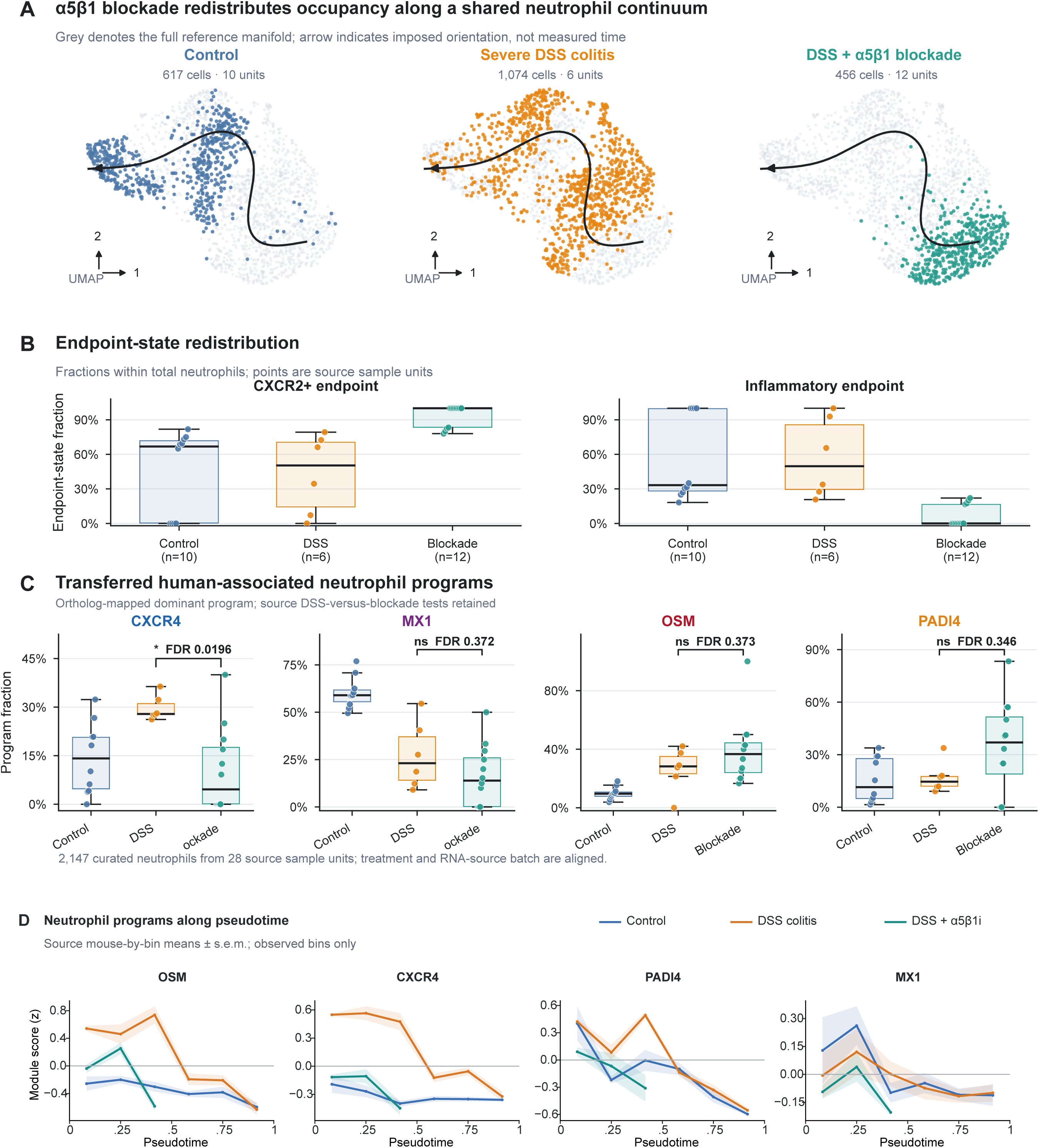
Neutrophil occupancy and transferred programs after ATN-161. (A) Condition occupancy and Slingshot curve rooted in the Cxcr2-associated state. (B) Fractions assigned to two endpoints. (C) Fractions assigned to transferred human neutrophil programs. Points indicate source units; boxes, median/interquartile range; whiskers, 1.5×interquartile range. DSS-versus-blockade Wilcoxon tests use BH correction across four programs. The inventory contains 2,147 cells and 28 units (10 control, 6 DSS, 12 blockade); Figure 5C excludes four control units upstream. (D) Program scores across pseudotime; 60 source-unit-by-bin means±SE. Treatment and batch are confounded; cross-sectional curves do not establish temporal transitions.

**Supplementary Figure 9.**
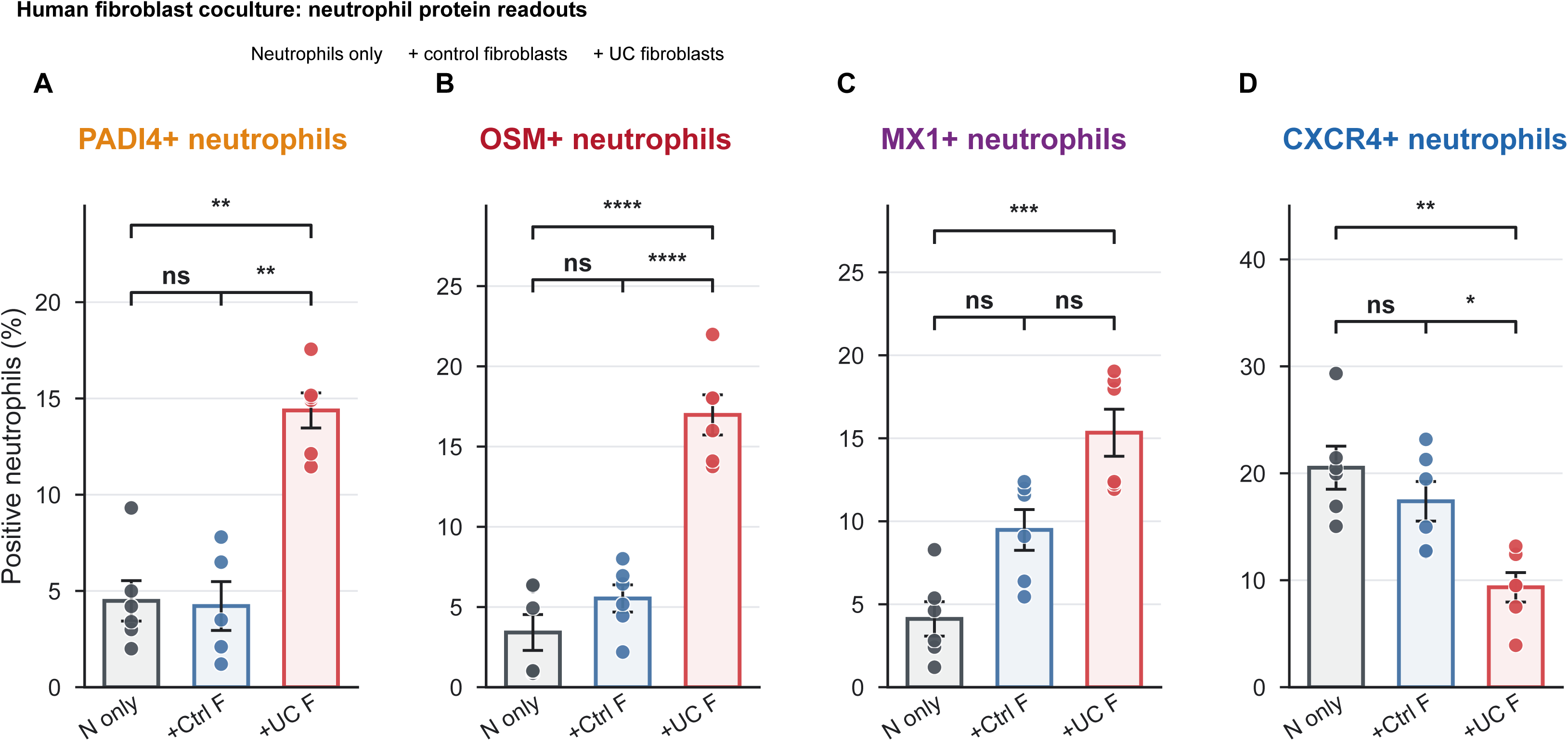
Marker-positive neutrophils in human fibroblast coculture. (A–D) PADI4-, OSM-, MX1-, and CXCR4-positive neutrophil percentages after culture alone, with control fibroblasts, or with UC fibroblasts. Source observations, mean±SEM designation, and significance annotations are retained. Samples and gates are not assumed to match Figure 3B,C. Marker-positive percentages do not establish mutually exclusive RNA states or effector activity.

**Supplementary Figure 10.**
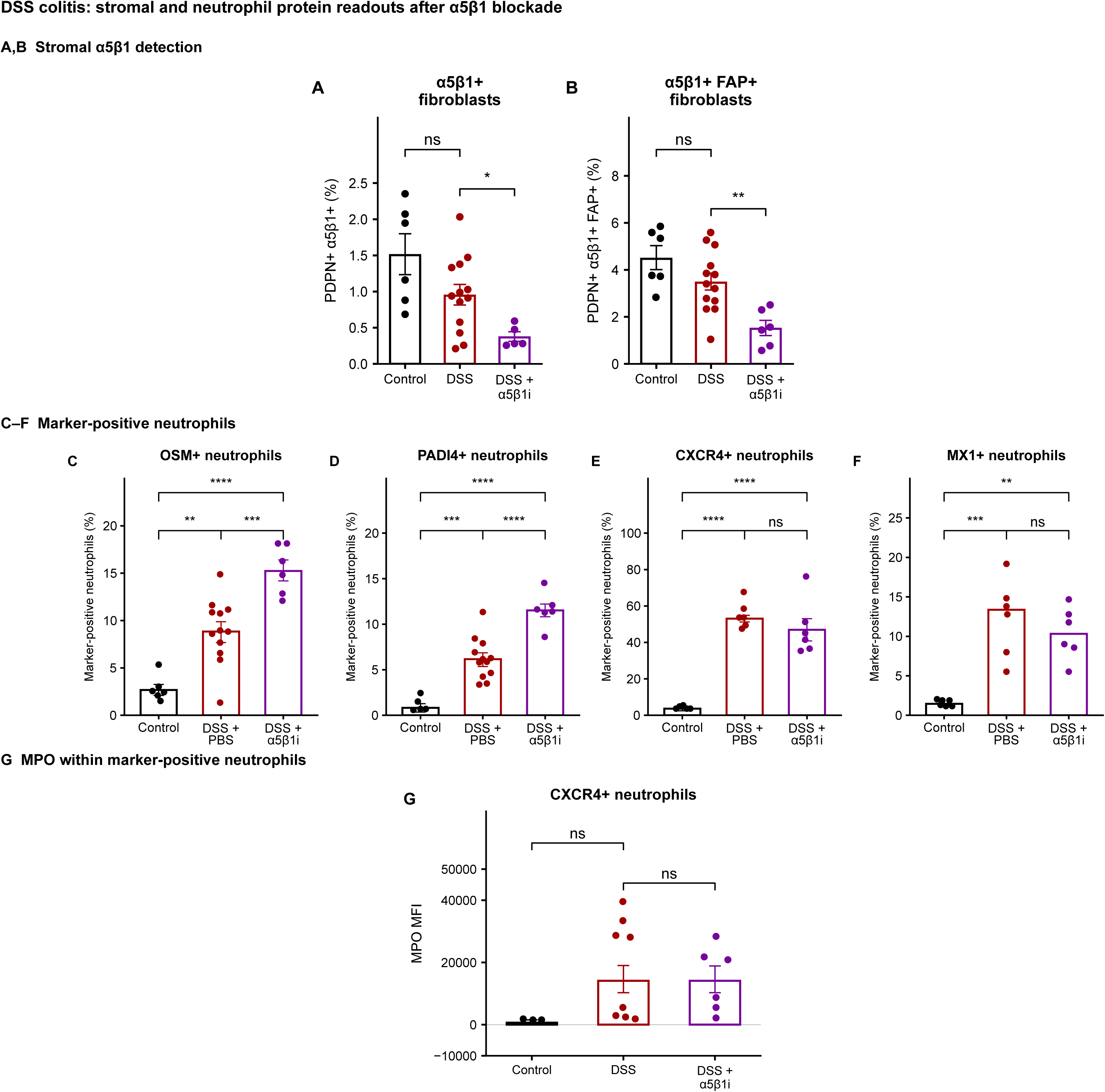
Stromal detection and complementary neutrophil protein measurements after ATN-161. (A,B) Detectable α5β1-positive fractions among PDPN-positive and PDPN-positive/FAP-positive fibroblasts. (C–F) OSM-, PADI4-, CXCR4- and MX1-positive percentages within neutrophils. (G) MPO fluorescence within CXCR4-positive neutrophils; OSM- and PADI4-subset MPO appear in Figure 5B. Black, control; dark red, DSS+PBS; purple, DSS+ATN-161. Source observations, error bars and annotations are retained; MX1 has no added error bars. Control-versus-DSS annotations in A,B,G retain reconstructed-point Welch/BH tests, with the three-endpoint MPO correction family shared with Figure 5B. Other annotations retain source tests. Lower integrin detection may reflect receptor occupancy or epitope interference. Marker-positive percentages do not establish absolute subset expansion or effector function.

**Supplementary Figure 11.**
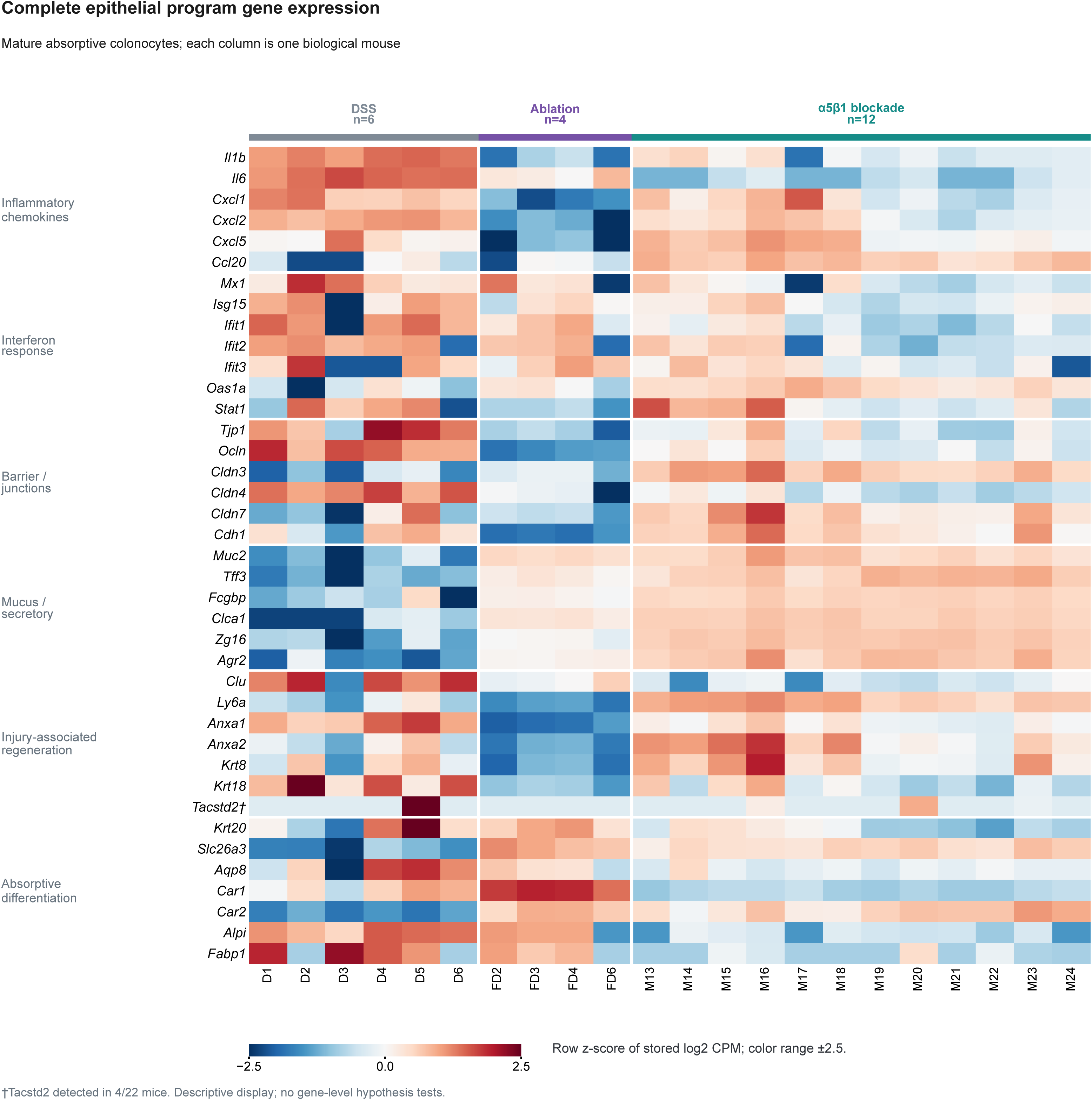
Complete epithelial program gene expression in mature absorptive colonocytes. Stored pseudobulk expression of all 39 genes from the six epithelial programs in mature absorptive colonocytes. Columns represent six DSS, four FAP-ablation and twelve α5β1-blockade biological mice, each with ≥20 cells in this state. Genes are grouped by program; mice are ordered by condition and identifier. Colors show gene-wise z-scores across the 22 equally weighted mice, calculated from existing log₂ counts per million using the population SD and saturated at ±2.5. White separators distinguish programs and conditions. All genes were detected in at least three mice; †Tacstd2 was detected in only 4/22. The original normalization includes a prior count. This descriptive display does not imply gene-level significance, protein abundance, epithelial lineage conversion or functional repair. Treatment and acquisition batch are confounded.

**Supplementary Figure 12.**
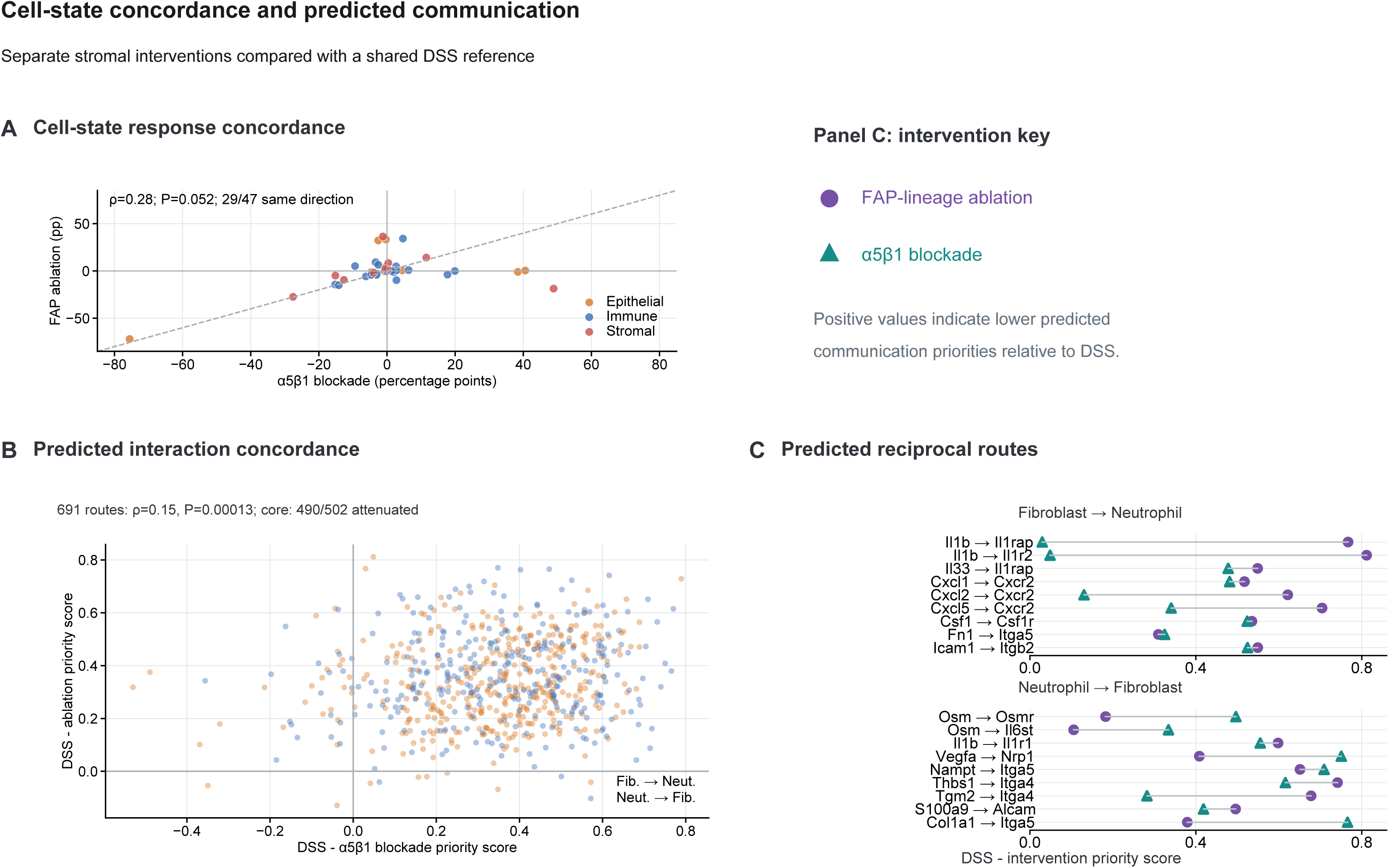
Cell-state concordance and predicted communication responses to stromal interventions. (A) Within-compartment proportion differences for 47 shared cell states; dashed line, equal differences; Spearman correlation describes concordance. These comparisons use six DSS, five ablation and twelve blockade biological mice. (B) Matched communication-priority contrasts for 691 routes; orange, fibroblast-to-neutrophil; blue, neutrophil-to-fibroblast. Of 502 jointly DSS-induced routes, 490 have lower inferred priority after both interventions. (C) Selected reciprocal-route differences; purple circles, ablation; teal triangles, blockade. Positive values indicate lower inferred priorities relative to DSS. Communication analyses retain their separate source eligibility and sparse-cell limitations, detailed in Supplementary Methods. Comparisons derive from separate experiments sharing a DSS reference; treatment and acquisition batch are confounded. Inferred priorities do not establish signaling activity or direct mechanisms.

## Supplementary Methods

### Study design and analytical units

The CODEX antibody panel is listed in Supplementary Table 1. Human tissue, CODEX, coculture, and DSS-colitis analyses used endpoint-specific eligibility (Supplementary Table 2). Patients, recorded culture samples or donors, and biological mice were the analysis units where established; cells were not independent replicates. Figure 3D comprises six independent experiments per group on the same scale with the same normalization. Flow cytometry, RNA, and elastase assays were not assumed to be matched. Collection of human blood and biopsies for coculture, scRNA-seq, and CODEX analyses were approved by Stanford University (IRB 28437, 60958, and 52317). Mouse procedures were approved by the Stanford University IACUC (22000 and 27715) and Mayo Clinic IACUC (A00008640-26). Experimental procedures are reported in the main Methods. References below are numbered independently of the main manuscript.

**Supplementary Table 1.** CODEX antibody panel and acquisition settings. The panel comprised 53 antibodies plus DAPI. Substance P was not used. Antibodies were conjugated to the listed oligonucleotide barcodes and visualized with the indicated fluorescent reporters.

| Marker | Oligo | Reporter | Cycle | Clone | Catalog no. | Manufacturer | Exposure (ms) | Dilution |
| --- | --- | --- | --- | --- | --- | --- | --- | --- |
| Vimentin | 7 | AF488 | 1 | RV202 | 550513 | BD Biosciences | 50 | 1:100 |
| BCL2 | 8 | AF488 | 2 | 124 | NBP2-50116 | Novus Biologicals | 150 | 1:100 |
| CD15 | 15 | AF488 | 3 | HI98 | 14-0159-82 | Invitrogen | 10 | 1:200 |
| CD194 | 21 | AF488 | 17 | Polyclonal | PA5-99885 | Invitrogen | 150 | 1:100 |
| CD19 | 28 | AF488 | 4 | FMC63 | NBP2-52716-0.2mg | Novus Biologicals | 90 | 1:50 |
| EpCAM | 41 | AF488 | 5 | Ber-EP4 | BE0386 | InVivoMab/ BioXCell | 300 | 1:50 |
| CD8 | 43 | AF488 | 7 | C8/144B | 90257SF | Cell Signaling Technology | 50 | 1:100 |
| αSMA | 44 | AF488 | 6 | 1A4/asm-1 | NBP2-33006-0.1mg | Novus Biologicals | 10 | 1:200 |
| Collagen IV | 46 | AF488 | 8 | 1043 | NBP1-97716 | Novus Biologicals | 150 | 1:100 |
| FcyRIα | 26 | AF488 | 9 | UMAB74 | LS-C796743-100 | LS Bio | 30 | 1:100 |
| CD11c | 67 | AF488 | 10 | EP1347Y | ab216655 | Abcam | 50 | 1:100 |
| CD127 | 66 | AF488 | 11 | BLR177J | NBP3-14749 | Novus Biologicals | 30 | 1:100 |
| CXCR4 | 68 | AF488 | 12 | Polyclonal | NB100-56437 | Novus Biologicals | 150 | 1:50 |
| PADI4 | 29 | AF488 | 13 | OTI4H5 | CF504813 | OriGene | 300 | 1:200 |
| CD68 | 72 | AF488 | 14 | KP1 | 916104 | BioLegend | 30 | 1:100 |
| Podoplanin | 32 | AF488 | 15 | NC-08 | 337002 | BioLegend | 150 | 1:50 |
| MX1 | 33 | AF488 | 16 | OTI2G12 | NBP2-72838 | Novus Biologicals | 150 | 1:200 |
| CD134 | 2 | ATTO550 | 1 | UMAB276 | LS-C799259-100 | LS Bio | 300 | 1:200 |
| CD56 | 5 | ATTO550 | 2 | MRQ-42 | 156R-200UG | Cell Marque | 300 | 1:50 |
| CD66b | 11 | ATTO550 | 3 | 6/40c | 392902 | BioLegend | 30 | 1:100 |
| CD274 | 14 | ATTO550 | — | E1L3N | 85164SF | Cell Signaling Technology | 150 | 1:100 |
| CD3 | 20 | ATTO550 | 6 | D7A6E | 24581SF | Cell Signaling Technology | 150 | 1:100 |
| FcεRIα | 24 | ATTO550 | 7 | 9E1 | ab54411 | Abcam | 300 | 1:200 |
| Vista | 30 | ATTO550 | 8 | D1L2G | 82119SF | Cell Signaling Technology | 150 | 1:100 |
| OSM | 36 | ATTO550 | 9 | 17001.31 | NB120-10842 | Novus Biologicals | 90 | 1:100 |
| CD11b | 38 | ATTO550 | 10 | Polyclonal | NB110-89474 | Novus Biologicals | 150 | 1:100 |
| FAP | 49 | ATTO550 | 11 | FAP/4853 | NBP3-14220 | Novus Biologicals | 150 | 1:50 |
| CD138 | 55 | ATTO550 | 12 | # 359103 | MAB2780 | R&D Systems | 150 | 1:100 |
| CD45 | 56 | ATTO550 | 13 | HI30 | NBP1-79127 | Novus Biologicals | 150 | 1:100 |
| α5β1 | 57 | ATTO550 | 14 | M200 (Volociximab) | ab275977 | Abcam | 90 | 1:100 |
| CK7 | 63 | ATTO550 | 15 | KRT7/760 | NBP2-47939-0.1mg | Novus Biologicals | 150 | 1:100 |
| LAG-3 | 71 | ATTO550 | 16 | D2G4O | 25848SF | Cell Signaling Technology | 300 | 1:50 |
| CD117 | 74 | ATTO550 | 17 | 2B8 | NB100-77477-0.1mg | Novus Biologicals | 150 | 1:100 |
| CD4 | 62 | ATTO550 | 4 | EPR6855 | ab181724 | Abcam | 150 | 1:100 |
| CD31 | 3 | Cy5 | 1 | C31.3 + C31.7 + C31.10 | NBP2-47785-0.1mg | Novus Biologicals | 20 | 1:200 |
| T-bet | 6 | Cy5 | 2 | 4B10 | NBP1-43298-0.1mg | Novus Biologicals | 300 | 1:200 |
| HLA-DR | 23 | Cy5 | 3 | TAL 1B5 | NB600-989 | Novus Biologicals | 6 | 1:200 |
| CD163 | 25 | Cy5 | 4 | ED2 | NBP2-39099-100ug | Novus Biologicals | 50 | 1:200 C00, 1:50 C02 |
| CD20 | 81 | Cy5 | 5 | L26 + IGEL/773 | NBP2-44747-0.1ml | Novus Biologicals | 150 | 1:200 |
| CD1a | 69 | Cy5 | 6 | O10+C1A/711 | NBP2-34698-0.1mg | Novus Biologicals | 90 | 1:100 |
| Ki67 | 42 | Cy5 | 7 | B56 | ab279657 | Abcam | 12 | 1:100 |
| P53 | 45 | Cy5 | 8 | PAb 240 | NB200-103 | Novus Biologicals | 150 | 1:100 |
| TCR$\gamma\delta$ | 53 | Cy5 | 9 | E2E9T | 61129SF | Cell Signaling Technology | 300 | 1:50 |
| HLA-ABC | 70 | Cy5 | 10 | EPR22172 | ab239788 | Abcam | 15 | 1:100 |
| CD16 | 52 | Cy5 | 11 | D1N9L | 72204SF | Cell Signaling Technology | 150 | 1:100 |
| CD140a | 59 | Cy5 | 12 | Polyclonal | AF-307-NA | Novus Biologicals | 150 | 1:200 |
| CD25 | 60 | Cy5 | 13 | Polyclonal | NBP2-38730 | Novus Biologicals | 150 | 1:200 |
| FOXP3 | 61 | Cy5 | 14 | 236A/E7 | 14-4777-82 | Invitrogen | 150 | 1:200 |
| CD14 | 76 | Cy5 | 15 | 1H5D8 | NBP2-37295 | Novus Biologicals | 90 | 1:100 |
| CD34 | 77 | Cy5 | 16 | QBEnd/10 | NBP2-32932-0.1mg | Novus Biologicals | 150 | 1:100 |
| CD279 | 79 | Cy5 | 17 | EH12.2H7 | MCA6133F | Bio-Rad | 150 | 1:200 |
| Chromogranin A | 75 | Cy5 | 18 | CGA/414 | NBP2-33198 | Novus Biologicals | 150 | 1:200 |
| TIGIT | 17 | Cy5 | 19 | TIGIT/3017 | NBP2-79927 | Novus Biologicals | 150 | 1:50 |

Oligo, oligonucleotide barcode identifier; ms, milliseconds. DAPI (BD Pharmingen, 564907) was used at 1:300. Run-specific dilution codes for CD194 and CD163 are retained from the assay worksheet. The $\alpha 5\beta 1$ reagent was clone M200 (Volociximab), rabbit IgG chimeric (Abcam, ab275977).

**Supplementary Table 2.**
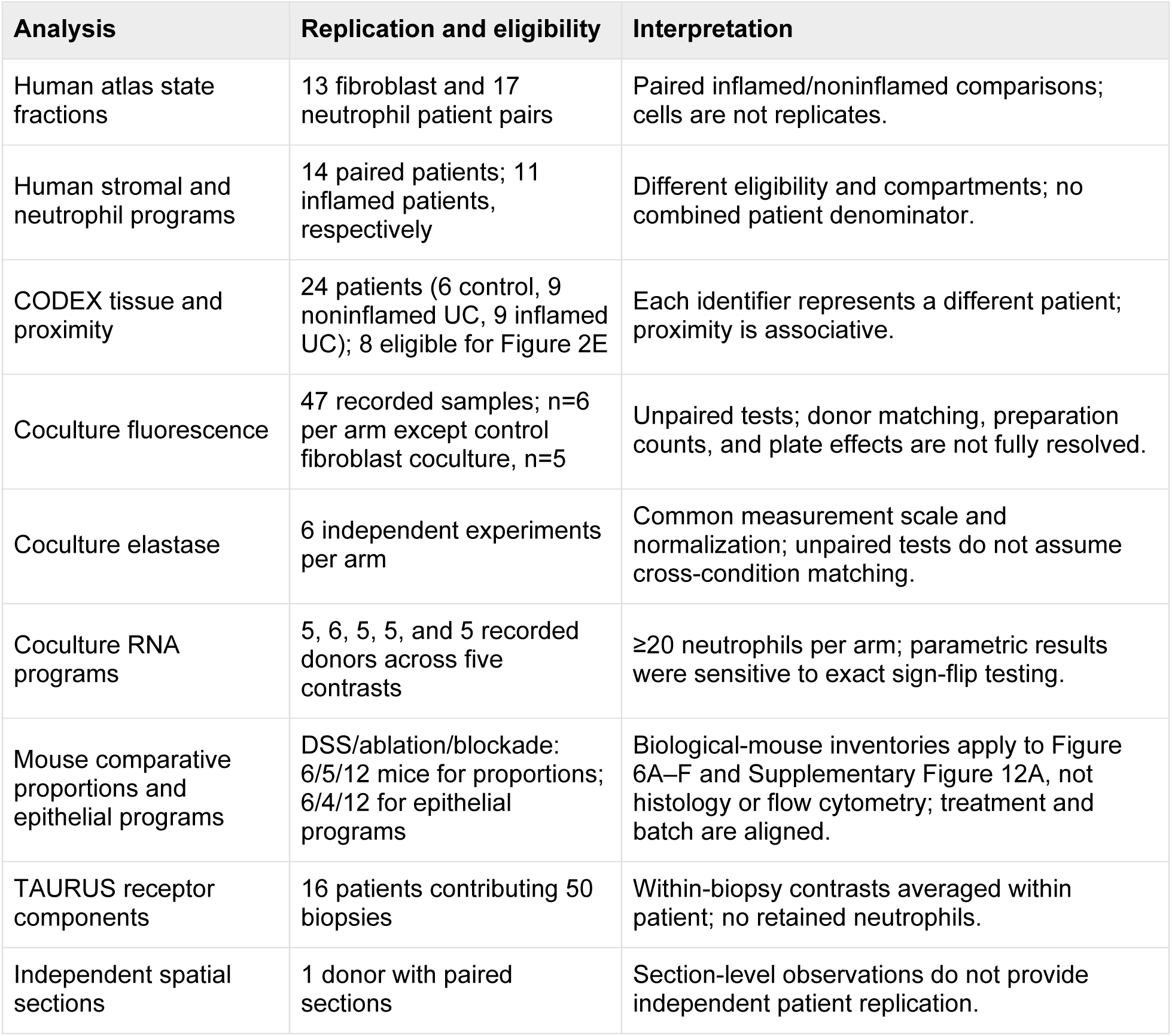
Analytical populations and interpretation.

### Discovery tissue analyses

Discovery analyses reused the published UC atlas and its annotations.^1^ The study is registered as SCP3755 in the Broad Institute Single Cell Portal.^2^ Focused displays contain 5,187 fibroblasts and 38,684 neutrophils; paired state analyses include 13 fibroblast and 17 neutrophil patient pairs, and receptor-expression analyses include 10 biopsies. Figure 1C evaluates fibroblast subtypes; Figure 1D evaluates the broader stromal compartment. Program scores use normalized RNA expression. Noninteger values in the available count layer limit reconstruction of the original count-processing history.

Paired stromal programs (Figure 1D; Supplementary Figure 1A). All Level 1 stromal cells were included, encompassing fibroblasts, smooth-muscle cells, and pericytes. Normalized expression was averaged across available genes per cell, retaining zeros, then across cells per patient–condition. Fourteen patients had paired samples. Paired Wilcoxon signed-rank tests used the asymptotic approximation with continuity correction and BH adjustment across eight programs. Open points mark <10 cells. Requiring ≥5, ≥10, or ≥20 cells in both samples retained 9, 6, or 5 pairs. The ≥5-cell analysis retained significant FAP-inflammatory/recruitment scores; no program remained BH-significant at ≥10 cells. These compartment-level means are sensitive to composition and sparse recovery.

The eight fibroblast-associated gene sets were FAP-inflammatory (FAP, PDPN, CXCL1, CXCL5, CXCL6, CXCL8, CSF3, IL6, ICAM1); α5β1 adhesion (ITGA5, ITGB1, FN1, PTK2, SRC, PXN, VCL, TLN1, ACTN1, RHOA, ROCK1, MYL9); neutrophil recruitment (CXCL1, CXCL2, CXCL3, CXCL5, CXCL6, CXCL8, CSF3); OSM response (OSMR, LIFR, IL6ST, STAT3, SOCS3, CEBPD, JUNB, FOSL2, CXCL1, CXCL8); matrix remodeling (COL1A1, COL1A2, COL3A1, COL5A1, COL6A1, FN1, POSTN, MMP2, MMP3, MMP14, TIMP1); TGFβ response (TGFBR1, TGFBR2, SMAD2, SMAD3, SERPINE1, CTGF, COL1A1, COL3A1); NFκB/AP1 (NFKBIA, NFKB1, RELA, TNFAIP3, JUN, JUNB, FOS, FOSB, ATF3); and YAP mechanotransduction (YAP1, WWTR1, TEAD1, CTGF, CYR61, ANKRD1, AMOTL2). Overlapping genes mean the modules are not biologically independent pathways.

Matched neutrophil programs (Figure 1E). The same 11 inflamed UC patients had ≥20 cells in each OSM-, CXCR4-, PADI4-, and MX1-associated state (27,621 cells). OSM, CXCR4, PADI4, and MX1 were removed wherever present in nine investigator-defined modules. Normalized expression was averaged across retained genes, within patient– state, and equally across patients. Heatmap values are z-scores across four state means within each program (population SD); tests use unscaled scores. Friedman omnibus tests use patient blocks and an asymptotic chi-square reference distribution, with BH correction across all nine modules, including three not displayed. These tests do not establish pairwise differences.

The nine original neutrophil gene sets were recruitment/migration (CXCR1, CXCR2, FPR1, FPR2, SELL, ITGAM, ITGB2); retention/aging (CXCR4, ICAM1, CD44, SELL, BCL2A1); granule/protease, originally Degranulation (MMP8, MMP9, ELANE, CTSG, AZU1, LTF, DEFA3, OLFM4); NET-associated (PADI4, MPO, ELANE, CTSG, HIST1H2AC, HIST1H4C); oxidative burst (CYBA, CYBB, NCF1, NCF2, NCF4, RAC2); inflammatory (OSM, IL1B, TNF, PTGS2, NFKBIA, CXCL8); interferon (MX1, ISG15, IFIT1, IFIT2, IFIT3, IFI6, OAS1); survival/immaturity (CSF3R, BCL2A1, OLFM4, LCN2, LTF, MPO, ELANE); and tissue injury (MMP8, MMP9, S100A8, S100A9, LCN2, CTSG, ELANE). Naming-gene exclusions apply to Figure 1E; gene coverage and retained definitions are supplied in source data. These discovery modules differ from the six coculture RNA programs in Figure 3E.

Sensitivity and annotation controls. Original scores were reproduced from normalized RNA. Minimums of 10, 20, and 50 cells in every state retained 12, 11, and 10 patients. Supplementary Figure 1B retains pooled, cell-weighted original scores across all conditions and seven annotations, including two erythrocytes, seven macrophages, and low-RNA cells. Values are standardized across annotations within each program; naming genes remain included. The nine non-neutrophil control cells are absent from the focused neutrophil expression object and Figure 1E. These controls do not independently validate state identities or effector function.

Paired coexpression (Supplementary Figure 1C). Nine patient pairs were compared by paired Wilcoxon tests. BH correction retains five routes: CSF3–CSF3R, CXCL1– CXCR2, IL1B–IL1R1, NAMPT–ITGA5/ITGB1, and OSM–OSMR. The NAMPT receptor score uses the lower ITGA5/ITGB1 expression. All original paired observations and statistics are retained; the three displayed q values are 0.055, 0.055, and 0.076. Coexpression scores differ from MultiNicheNet priorities and do not establish ligand binding or receptor assembly.

### Human participants and ethics

Analysis-specific human denominators reflect lineage recovery and eligibility; no combined denominator is assigned across datasets. CODEX identifiers P01–P24 represent 24 different patients. The published atlas describes its original recruitment and ethics procedures.^1^

### Transcriptomic processing and culture annotation

Culture annotation used DecontX correction,^3^ fibroblast-specific principal-component analysis, donor Harmony integration,^4^ and Leiden clustering.^5^ The 238-cell audit included low-confidence immune-like assignments; 170 fibroblasts supported communication analyses. The neutrophil audit used scVI–Harmony embeddings^4,6^ and excluded low-signal and contaminant-like cells from resolved-state program analyses. Audit and cleanup counts represent different processing stages. Donor-blocked edgeR quasi-likelihood models were retained where documented.^7^ Recorded labels do not independently establish biological pairing.

### CODEX measurements and neighborhoods

Tissue sections were baked at 70°C for 60 minutes, deparaffinized in xylene, and rehydrated through ethanol. Heat-induced epitope retrieval was performed at pH 9 and 97°C for 10 minutes using a PT Module. Sections were blocked and incubated overnight at 4°C with a 53-antibody panel (Supplementary Table 1) and DAPI in 300 µL of blocking solution. The α5β1-specific antibody was clone M200 (Volociximab; rabbit IgG chimeric; Abcam, ab275977), used at the recorded 1:100 dilution. Other key antibodies included FAP (FAP/4853), CD66b (6/40c), CD15 (HI98), CD16 (D1N9L), OSM(17001.31), CXCR4 (polyclonal), PADI4 (OTI4H5) and MX1 (OTI2G12). After staining, sections were fixed using 1.6% paraformaldehyde, ice-cold 100% methanol, and an additional fixative reagent. Immediately before imaging, slides were fitted with a flow cell and equilibrated in running buffer. Fluorescent oligonucleotide reporters complementary to the antibody barcodes were prepared in a 96-well plate containing running buffer, nuclear stain, and assay reagent. Multicycle imaging was performed using PhenoImager Fusion software (version 2.2) with cycle-specific acquisition parameters. Image processing generated registered, background-subtracted, multichannel whole-slide images for downstream spatial analyses.

### CODEX preprocessing and Seurat clustering

Per-cell mean fluorescence, nuclear mean fluorescence and cell-centroid coordinates were exported from QuPath.^8^ Cell-level intensity profiles were stored in the Akoya assay of a Seurat object (Seurat 5.5.0; SeuratObject 5.4.0; R 4.5.3).^9^ The combined object contained 245,020 cells from 24 patients. NormalizeData used centered-log-ratio normalization with margin=2; ScaleData centered and scaled features, and FindVariableFeatures used the vst method. Principal-component analysis retained 30 components (seed 42). Harmony 2.0.5 adjusted these components by PatientID.^4^ A shared-nearest-neighbor graph was built from Harmony dimensions 1–10 with k=20, Euclidean distance and SNN pruning at 1/15. Louvain clustering used FindClusters at resolution 1.0 (algorithm 1; 10 starts; 10 iterations; seed 0).

The cohort UMAP used Harmony dimensions 1–30, cosine distance, 30 neighbors, minimum distance 0.3, 200 epochs and seed 42. The stored normalization, scaling and PCA included 54 feature columns, including a legacy Substance P column. Substance P was not used in the staining panel and is excluded from Supplementary Table 1; its retained column in the clustering object should not be interpreted as evidence of measured Substance P staining.

### CODEX cell annotation and atlas-guided phenotypes

Cell annotation combined available lineage-protein profiles with selected marker relationships and neutrophil-state definitions from the published UC single-cell atlas.^1^ Epithelial and enteroendocrine annotations were supported by EpCAM/CK7 and chromogranin A, respectively; endothelial profiles were evaluated with CD31. CD140a (PDGFRA), vimentin, podoplanin, collagen IV, αSMA and FAP provided stromal context. CD45 and lineage markers distinguished immune populations: CD3 with CD4 or CD8 for T cells; FOXP3 and CD25 for regulatory T cells; CD19/CD20 for B cells; CD138 for plasma cells; CD68, CD163 and CD14 for macrophages; and CD11c, HLA-DR and CD1a for dendritic-cell profiles. Labels were based on combinations of markers rather than any single marker alone.

Neutrophil identity was supported by CD66b, CD15, CD16 and CD11b, with OSM, CXCR4, PADI4 and MX1 used to distinguish atlas-informed protein phenotypes. The final annotations retained the existing resolution-0.8 reference labels by cell identifier for 226,659 cells from 23 patients. For P07, labels were assigned to 18,361 cells by a 25-nearest-neighbor majority vote in global Harmony dimensions 1–10; median vote support was 0.92. This transfer was between CODEX cells, rather than direct computational transfer from the RNA atlas. The final set contained 12 broad lineages and 14,906 neutrophils, including OSM-, CXCR4-, PADI4- and MX1-associated groups and a residual unpolarized group (Supplementary Figure 2). These protein phenotypes provide an atlas-informed interpretation and do not establish one-to-one equivalence with transcriptomic states or distinct effector functions.

Spatial analyses used the retained cell labels, centroids and recorded marker intensities. Neutrophil marker summaries used the exported nuclear mean-fluorescence fields, whereas FAP and α5β1 fibroblast thresholds used the recorded cell-level raw or normalized intensities specified below. These intensity summaries are operational assay readouts and do not by themselves establish subcellular localization.

### Cellular neighborhoods and proximity analyses

Six neighborhoods summarize cell-type composition within 50 μm with equal patient weighting. Exact patient-level occupancy comparisons use BH correction across 36 tissue and neutrophil tests. Tissue occupancy is the fraction of assigned cells, not absolute density.

Availability-adjusted proximity (Figure 2D). Pooled-UC centered-log-ratio upper-quartile thresholds were 2.0520675548890175 for FAP and 0.9314878591724798 for α5β1. Marker categories were permuted within connected-component, epithelial-distance, and local-density strata, preserving fibroblast positions and joint category abundance. Eligibility required ≥10 reference fibroblasts, ≥5 expected edges, and ≥10 eligible neutrophils. Patient values were log₂[(observed+0.5)/(expected+0.5)]; group means use t CIs. BH correction covers 24 planned null/group comparisons.

Continuous proximity (Figure 2E). A two-Gaussian mixture fitted to log1p raw fibroblast α5β1 signal gave equal total weight to each patient. Posterior ≥0.50 corresponded to raw signal ≥35.647672463540204 and selected 806/44,516 fibroblasts. The fitted density was unimodal; this operational high-signal gate differs from Figure 2D quartiles. FAP-high and α5β1-positive populations can overlap. Epithelial/endothelial references use annotations; CD31 was present in the clustering object but absent from the downstream proximity-analysis export.

Exposed neutrophils were summarized in 150-μm blocks with ≥3 neutrophils. Models required ≥30 exposed neutrophils, ≥12 blocks, and design-matrix condition number <30. Standardized block-mean log1p marker expression was regressed jointly on negative standardized log1p reference-cell distances, fibroblast count, and total density. Distances had no maximum-radius or additional connected-component restriction. Eight inflamed patients qualified (P07–P12, P14, P15); some had only 1–4 α5β1-positive fibroblasts. Equally weighted patient coefficients use one-sample t tests, pointwise t CIs, and BH correction across 24 outcomes. Gate uncertainty was not propagated.

Supplementary spatial controls use different reference definitions. Within-patient immune-lineage label exchangeability generated expected edges in Supplementary Figure 3A; 49 percentile combinations assess threshold sensitivity in B. The 17-patient model adjusts for density and anatomical proxies and uses 499 restricted-null permutations. These do not retest the Figure 2E gate.

### Coculture conditions and FACS

Fresh control-blood neutrophils were cocultured for 16 hours with fibroblasts isolated separately from pooled control or Mayo-2 UC biopsies, using the main Methods protocol. ATN-161 (20 μM), FK-866 (10 μM), and dual-treatment arms were retained. Inhibitor-only and coculture effects were analyzed separately; the initial fibroblast seeding density is not an endpoint coculture ratio. Independent fibroblast preparation counts, pooling across versus within patients, and culture matching remain unresolved. No extracellular NAMPT-specific neutralization or drug synergy is inferred.

Representative flow samples minimize the mean squared standardized distance between five marker medians and their within-arm medians: P1-6, alone; P5-2, control fibroblasts; P3-5, UC fibroblasts; P4-2, UC fibroblasts+ATN-161. Each sample supplies all five histograms. Scatter/singlet/CD16/CD11b gates retain CXCR4-negative cells. Histograms use arcsinh(fluorescence/150), common within-marker bins, pooled 0.1st– 99.9th percentile limits, and 2.5% padding; ticks show untransformed values. No smoothing or event subsampling was applied.

Figure 3C compares sample-median compensated fluorescence scaled by marker SD across 47 samples (n=6 per arm except control coculture, n=5). Nine contrasts across five markers form 45 exact unpaired permutation tests with BH correction. Negative compensated values are retained. MX1/PADI4 channel assignments conflict with protocol annotations; most files lack viability measurements, and treatment may be confounded with plate. These tests do not assign drug action to fibroblast α5β1.

### Elastase and additional protein assays

NET-associated elastase activity was measured after removal of unbound elastase and S7-nuclease digestion using Cayman kit 601010, with chromogenic detection at 405 nm. Figure 3D includes six independent experiments per arm, measured on the same scale using the same normalization (mean±SD). Each exact unpaired six-versus-six test enumerates 924 allocations; Holm correction covers three adjacent comparisons. The first three arms were reconstructed from source artwork and their plotted values were author-verified; the ATN-161 observations were supplied numerically (12, 18, 16, 14, 14, 16; mean 15.0, SD 2.10). The unpaired analysis does not assume cross-condition donor matching. This activity endpoint alone does not establish NET formation.

Additional protein assays retain author-verified plotted observations and assay-specific annotations. Verification of plotted values does not resolve antibody identity, biological independence or missing test metadata. Figure 5B presents MPO fluorescence within OSM-positive and PADI4-positive neutrophils; Supplementary Figure 10G retains the CXCR4-positive gate. Marker-positive percentages use all neutrophils as the denominator (Supplementary Figure 10C–F), whereas MPO is measured within each indicated positive gate; the two endpoints are not interchangeable. Control-versus-DSS ns labels derive from reconstructed-point Welch tests, with BH correction across two stromal endpoints (Supplementary Figure 10A,B) or the original three MPO endpoints (Figure 5B and Supplementary Figure 10G). Moving the two MPO plots into the main figure does not change that correction family. DSS-versus-blockade MPO annotations retain source tests. Supplementary Figure 10C–F annotations come from the latest supplied protein measurements; MX1 has no added error bars. Supplementary Figures 7C–F and 9 retain their original summary/error-bar and significance designations without a common recomputed testing family.

### Neutrophil RNA programs and predicted ligand support

Six investigator-curated culture programs were scored in resolved neutrophils from recorded donor–condition arms with ≥20 cells. Expression was log1p(count/total UMI×10,000); reference means/variances equally weighted untreated donor–condition groups. Reference SD<10⁻⁸ was replaced by one. Standardized gene scores were averaged within programs and samples. Paired t tests and 95% t CIs use n−1 degrees of freedom; the five contrasts have n=5,6,5,5,5 recorded donors. BH correction covers six programs×five contrasts (30 tests), yielding 16 discoveries. Exact paired sign-flip tests yield no BH discoveries across the same family. Modules overlap and do not measure functional capacity.

The complete investigator-curated module membership is retained in figure source data. Chemotaxis comprises CXCR1, CXCR2, CXCR4, FPR1, FPR2, C5AR1, CCR1 and CCRL2. Granule-associated genes comprise LTF, BPI, CAMP, LCN2, MMP8, MMP9, OLFM4, CEACAM8 and FCGR3B. Hypoxia/glycolysis comprises HIF1A, SLC2A1, HK2, PFKP, ALDOA, GAPDH, PGK1, LDHA and ENO1. The inflammatory module comprises OSM, IL1B, TNF, NFKBIA, CXCL8, CCL3, CCL4, IL1R2 and SOCS3. The oxidase-associated module comprises CYBB, CYBA, NCF1, NCF2, NCF4, RAC2 and MPO. The interferon module comprises MX1, ISG15, IFIT1, IFIT2, IFIT3, IFI6, OAS1, OAS2, OASL, RSAD2, IRF7 and STAT1. These definitions are not independent gene-level differential-expression claims.

RNA composition analyses use four-state centered-log-ratio contrasts with donor-specific variance. The α5β1-by-context interaction subtracts the inhibitor effect without fibroblasts from its effect with UC fibroblasts. Supplementary Figure 6 includes six recorded donors with no positive minimum-cell filter; none of four state interactions passes correction. These analyses differ from program scores, marker-positive gates, and fluorescence medians.

Figure 3F displays six routes selected for opposite exposure/blockade score directions, not a network-wide reversal test. Figure 3G uses the union of genes tested in resolved-neutrophil pseudobulk analysis and present in the NicheNet ligand–target prior.^10^ Within-ligand midrank percentiles retain zero weights and tied ranks. Color is mean percentile across available module genes; dot area is 25+4×the count at percentile ≥0.90, printed within the dot. Zero is an observed count. This prior-based summary has no enrichment test or inferred activation direction.

### Mouse experiments and histology

Mouse procedures and approvals are reported in the main Methods. The ablation RNA inventory includes five control, six DSS, and five ablation biological mice plus four pseudo-derived ablation units. These four units remain in the original Milo model but are excluded from Figure 6A–F and Supplementary Figure 12A. The blockade main RNA inventory in Figure 5C excludes C7–C10, leaving 24 units (6/6/12); Supplementary Figure 8 retains 28 (10/6/12). These RNA inventories do not establish histology or flow-cytometry sample sizes.

Histology was graded by a blinded pathologist using the cited inflammation-scoring framework.^11^ Middle DSS fields received bounded luminance sharpening (Gaussian radius 2.0 pixels; amount 2.0; threshold 2/255; maximum correction 20/255), without generative image processing. Original crops and aspect ratios were retained. Displayed 100-μm bars use provisional historical lengths (23% of panel width); acquisition calibration and field identities remain unverified.

Milo summaries in Figures 4C and 5D retain FDR<0.05 calls, log-fold-change direction, and dominant neighborhood annotations.^12^ Overlapping neighborhoods are not independent state tests, and Osm/Padi4/Mx1/Cxcr4 RNA labels are not flow gates. The trajectory inventory contains two residual ablation neutrophils; the separate communication inventory contains four across three eligible units, including one pseudo-derived unit. These inventories are not interchangeable.

### Comparative epithelial and communication analyses

Figure 6C,D proportions use six shared DSS, five ablation and twelve blockade biological mice. Mean differences and pointwise percentile CIs use 10,000 mouse bootstraps. Mann–Whitney comparisons are BH-adjusted across all Level 3 states per intervention. Supplementary Figure 12A shows concordance of within-compartment percentage-point effects for 47 states. Recovered proportions do not establish absolute abundance.

Epithelial counts were aggregated by biological mouse, TMM-normalized,^13^ and scored as log₂ CPM (prior.count=1). Six programs contain 39 genes detected in the assay and with positive counts in ≥3 eligible mice. The ≥20-cell requirement retains 6 DSS, 4 ablation and 12 blockade mice; FD5 (16 cells) is excluded, not assigned zero. BH correction covers 12 whole-compartment tests (Figure 6A). Within-state analyses require ≥20 cells per mouse and ≥3 mice per group, with BH correction across 30 tests (Figure 6B). Transitional cells have n=6/3/unsupported; mature absorptive cells, 6/4/12; IFN/MHC-II-responsive cells, 6/4/5 (DSS/ablation/blockade). Rank-test P values and bootstrap mean-difference CIs concern different quantities.

Figure 6E displays stored mature absorptive-colonocyte pseudobulk expression for 16 biologically selected genes, with the selection fixed before inspecting their expression in these mice: Cxcl1, Cxcl2, Cxcl5, Ccl20, Muc2, Tff3, Fcgbp, Tjp1, Ocln, Cldn7, Slc26a3, Aqp8, Alpi, Clu, Ly6a and Tacstd2. Supplementary Figure 11 displays all 39 genes from the original six programs. Both displays retain the same 22 eligible mice (6/4/12); each contributes at least 20 mature absorptive colonocytes. Gene-wise z-scores use the mean and population SD across the 22 equally weighted mice; colors saturate at ±2.5, with uncapped scores and stored log₂ CPM values retained in source data. Mice are ordered by condition and identifier, without expression clustering. Tacstd2 is detected in 4/22 mice and is flagged; normalization retains the original prior count. These displays do not supply gene-level significance tests.

The original epithelial program membership is: Inflammatory chemokines (Il1b, Il6, Cxcl1, Cxcl2, Cxcl5, Ccl20); Interferon response (Mx1, Isg15, Ifit1, Ifit2, Ifit3, Oas1a, Stat1); Barrier / junctions (Tjp1, Ocln, Cldn3, Cldn4, Cldn7, Cdh1); Mucus / secretory (Muc2, Tff3, Fcgbp, Clca1, Zg16, Agr2); Injury-associated regeneration (Clu, Ly6a, Anxa1, Anxa2, Krt8, Krt18, Tacstd2); Absorptive differentiation (Krt20, Slc26a3, Aqp8, Car1, Car2, Alpi, Fabp1).

Figure 6F uses the existing whole-epithelium inflammatory chemokine, absorptive differentiation and mucus/secretory scores joined by biological mouse identifier. Both plots retain all 22 mice without jitter; small symbols denote mice and large crosses denote arithmetic group means. No normalization, gene programs, correlations, regression models, trajectories or hypothesis tests were added. The gene and individual-mouse panels are descriptive views of the existing expression and program-score data.

MultiNicheNet version 2.1.0^14^ combines expression, specificity, prevalence, differential expression, and predicted ligand activity. In Supplementary Figure 12B,C, across four contrasts, 691 matched nonambient routes were evaluable; 490 of 502 jointly DSS-induced routes had lower priority after both interventions. Scores are not P values or signaling flux. The blockade interaction comparison retains three mice, two with only 2 and 11 fibroblasts. DSS, ablation, and blockade source batches are M1, M3, and I1; treatment cannot be separated from batch.

### Independent human analyses

TAURUS^15,16^ (Zenodo version 3, published October 30, 2024) includes 22 UC patients and 108 biopsies before filtering (6 remission, 14 nonremission, 2 unavailable outcomes). Counts were log1p-normalized to 10,000 per cell; fixed programs were scored without remission labels. Pretreatment inflammation comparisons are paired within patient. Figure 7A sign effects equal (positive−negative differences)/nonzero differences, with 1,800 patient bootstraps and BH correction across six features; feature-specific n=9,8,8,8,8,4.

Activated fibroblasts use the THY1pos FAPpos PDPNpos annotation, excluding pericytes. Figure 7C requires ≥5 activated and ≥20 other fibroblasts per biopsy; 16 patients contribute 50 pre/post-treatment biopsies. Within-biopsy expression/codetection differences are averaged within patient, with 10,000 bootstrap resamples and exact paired sign-flip tests; BH correction covers four genes and two receptor-component pairs. Codetection depends on depth/dropout and does not establish assembled surface receptors. Site-matched trajectories include 6 remission and 13 nonremission patients. TAURUS retains no neutrophils; the myeloid-feedback score is a monocyte proxy.

SCP3818^17^ provides paired sections from one UC donor. Activated-FAP cells detect FAP and PDPN or THY1 within the stromal annotation (n=146); neutrophil-like myeloid cells detect ≥2 of CSF3R, CXCR1, CXCR2, FCGR3B, CEACAM8, and MPO (n=879). Figure 7E uses 1,000 size-matched stromal sets. Figure 7F additionally matches epithelial-neighbor fraction among 30 nearest cells and normalizes distances by median all-cell nearest-neighbor spacing. Its pointwise null envelope is not a patient-level CI or validated histologic-compartment adjustment.

Figure 7G retains 17 normalized enrichment scores across seven pathways. Human contrasts use OSM-like α5β1-by-UC interactions except oxidative phosphorylation, which uses inhibition within UC coculture; mouse scores use total-neutrophil pseudobulk. Four unavailable combinations are NA. Different populations, gene universes, and contrasts preclude pooled replication claims.

### Statistical reporting

Tests were two-sided with endpoint-specific correction families; intervals are pointwise. Biological units were resampled where established. Nonsignificance does not establish equivalence, proportions do not establish absolute abundance, and transcript/protein measurements do not establish effector function. Original assay annotations are identified where full test metadata are unavailable.

### RNA module definitions and trajectory interpretation

Slingshot^18^ uses a curated Cxcr2-associated root and cross-sectional cells. Transferred human neutrophil programs use mapped mouse genes; fractions retain BH correction across four programs. Pseudotime displays source-unit-by-bin means at supported bins only, without lineage tracing or formal treatment-by-curve tests.

### Additional gene sets and target examples

Transferred mouse programs include genes beyond their naming markers. CXCR4-associated examples are Cxcr4, Hilpda, Plin2, Vegfa, Hmox1, Bhlhe40, and Sqstm1; OSM-associated examples are Osm, Il1b, Socs3, Nfkbia, Cxcl2, and Pfkfb3. The PADI4-associated program includes granule and transcriptional-regulatory genes. Predicted fibroblast target examples include Cxcl1/Vcam1/Cd44/Nfkb1 for OSM in ablation and Ccl2/Ccl7/Csf1/Cxcl10/Cxcl5 for IL1 in blockade. Target membership and pathway enrichment do not establish cell death, epithelial conversion, or receptor-specific mechanisms.

### Data and code availability

The discovery atlas is available at Broad Single Cell Portal SCP3755;^2^ TAURUS is deposited in Zenodo (version 3, 10.5281/zenodo.14007626);^16^ and the independent Xenium study is SCP3818.^17^ Scripts, selected derived tables, and main and supplementary figures are public at https://github.com/GubatanLab/UC-fibroblast-neutrophil-manuscript and archived as release 1.0.0 in Zenodo (DOI: 10.5281/zenodo.22881923). Underlying CODEX, human fibroblast–neutrophil coculture, and mouse colitis scRNA-seq datasets are available upon request from the corresponding author; they are not included in the public deposit.

Author names in bold designate shared co-first authorship

